# Severity of SARS-CoV-2 infection as a function of the interferon landscape across the respiratory tract of COVID-19 patients

**DOI:** 10.1101/2021.03.30.437173

**Authors:** Benedetta Sposito, Achille Broggi, Laura Pandolfi, Stefania Crotta, Roberto Ferrarese, Sofia Sisti, Nicola Clementi, Alessandro Ambrosi, Enju Liu, Vanessa Frangipane, Laura Saracino, Laura Marongiu, Fabio A Facchini, Andrea Bottazzi, Tommaso Fossali, Riccardo Colombo, Massimo Clementi, Elena Tagliabue, Antonio E Pontiroli, Federica Meloni, Andreas Wack, Nicasio Mancini, Ivan Zanoni

**Author notes:** These authors contributed equally (Benedetta Sposito, Achille Broggi). These authors contributed equally (Nicasio Mancini, Ivan Zanoni).

## Abstract

The COVID-19 outbreak driven by SARS-CoV-2 has caused more than 2.5 million deaths globally, with the most severe cases characterized by over-exuberant production of immune-mediators, the nature of which is not fully understood. Interferons of the type I (IFN-I) or type III (IFN-III) families are potent antivirals, but their role in COVID-19 remains debated. Our analysis of gene and protein expression along the respiratory tract shows that IFNs, especially IFN-III, are over-represented in the lower airways of patients with severe COVID-19, while high levels of IFN-III, and to a lesser extent IFN-I, characterize the upper airways of patients with high viral burden but reduced disease risk or severity; also, IFN expression varies with abundance of the cell types that produce them. Our data point to a dynamic process of inter- and intra-family production of IFNs in COVID-19, and suggest that IFNs play opposing roles at distinct anatomical sites.

## Introduction

Since the outbreak of the coronavirus disease 2019 (COVID-19) in early 2020, the novel, severe acute respiratory syndrome coronavirus 2 (SARS-CoV-2) has infected over 110 million people globally and caused more than 2.5 million deaths. SARS-CoV-2 infection can lead to acute respiratory distress syndrome (ARDS), with the most severe cases characterized by the presence of enhanced levels of pro-inflammatory cytokines in the bloodstream (*1–4*). Pattern recognition receptors (PRRs) of innate immune cells allow recognition of exogenous or endogenous molecules that are produced during microbial encounter, along with activation of antigen-specific adaptive immune responses aimed at eliminating the invading pathogen (*5*). Mouse models and retrospective human studies suggest that severity and death following SARS-CoV-2 encounter is correlated with exaggerated inflammation rather than viral load (*1–4, 6–8*). Consistent with this immuno-pathological etiology of severe COVID-19 is the efficacy of corticosteroids in treating hospitalized patients (*9*). Nevertheless, how a balance between the benefits (restricting viral replication and spread) and risks (inducing a cytokine storm) of efficient immune cell activation is achieved during COVID-19 remains a mystery.

Of the many inflammatory mediators produced upon infection with SARS-CoV-2, interferons (IFNs) have attracted much attention since the inception of the pandemic. IFNs belong to three major families: IFN-I (mainly represented by IFN-αs and IFN-β), IFN-II (IFN-γ), and IFN-III (IFN-λ1-4) (the most recent addition) (*10–12*). Upregulation of IFN-II in patients with severe COVID-19 (*1, 7*) is associated with increased PANoptosis, which exacerbates pathology and death (*7*). In contrast, the roles of IFN-I and IFN-III during SARS-CoV-2 infection have been a matter of debate. Indeed, IFN-I and IFN-III are among the most potent natural antivirals produced by mammals (*13, 14*) and share a large part of the signaling cascade that is downstream of their receptors (IFNAR and IFNLR, respectively). IFN-I and IFN-III also induce transcription of an overlapping pool of IFN-stimulated genes (ISGs). As such, the transcriptional signature of these two types of IFNs is almost indistinguishable (*14*). But given that expression of the IFNLR is restricted to just a few cell types (mainly epithelial cells and neutrophils), IFN-III can induce an antiviral state while simultaneously limiting inflammation-driven tissue damage (*14*). A seminal study showed that SARS-CoV-2, compared to other viruses, boosts the production of inflammatory mediators, while dampening the induction of ISGs and viral control in COVID-19 patients (*15*). Subsequent studies have confirmed that levels of IFN-I are highly impaired in the peripheral blood of patients with severe COVID-19 (*16*); also that such patients exhibit a profound suppression of IFN signaling (*17*) and a diminished and delayed production of IFN-III and IFN-I (*18*) compared to flu-infected patients. Nevertheless, regulation of IFN-I and IFN-III production following infection with SARS-CoV-2 appears to be more complicated. In fact, analysis of the bronchoalveolar lavage fluid (BALF) derived from the lower airways of COVID-19 patients has revealed a heightened immune response and increased ISG production (*19*). ISGs are also potently induced in peripheral blood monocytes of patients with severe COVID-19 (*2*), and production of IFNs is prolonged in the blood of a longitudinal cohort of COVID-19 patients (*1*). These discrepancies may, in part, be explained by the complexity and diversity of the patient cohorts, by the fact that ISGs rather than IFNs were measured, and by the heterogeneity of samples analyzed (e.g.: BALF vs. plasma).

Aside from the challenge of understanding if, when, and where IFNs are produced in patients infected with SARS-CoV-2, a major unanswered question is whether IFNs serve a protective or a detrimental function in COVID-19. Recent studies show that up to 15% of patients with severe COVID-19 have inborn errors of IFN-I immunity and autoantibodies against IFN-I (*20, 21*). Also, lack of an IFN signature is associated with a poor prognosis (*22*). These data strongly suggest that an efficient IFN response is essential for protecting against the development of a severe pathology. Other studies, however, report that dysregulated and prolonged production of IFNs in patients infected with SARS-CoV-2 is correlated with negative clinical outcomes (*1, 2*). We have also recently demonstrated that, after prolonged exposure to a respiratory virus or to viral ligands, the production of IFN-III, and to a lesser extent of IFN-I, impairs lung function and may trigger a severe pathology (*23, 24*). Thus, it is urgent to fully unravel the role of IFNs in the pathogenesis of COVID-19.

To define how IFN production varies at different anatomical sites during the progression of COVID-19, here we have analyzed a cohort of more than 250 subjects, including: i) SARS-CoV-2-infected patients with different clinical severity; ii) individuals with ARDS that was not driven by SARS-CoV-2 (either diagnosed with H1N1 influenza A virus or not); iii) patients with non-infectious lung pathologies; and iv) healthy controls. We analyzed the pattern and level of expression of IFNs and of pro-inflammatory cytokines in the upper or lower respiratory tract. Our data demonstrate that specific members amongst and between IFN families are differentially regulated, based on the location along the respiratory tract, the age of the patient, the severity of the pathology, and the viral load. Patients that were severely infected with SARS-CoV-2 displayed a unique inflammatory signature in the lung compared to that in the blood, or to the one found in patients with other types of ARDS. Finally, the production of specific members of the type I and type III IFN families was related to either immune cells or epithelial cells, and to the unique ability of specific PRRs to induce IFN expression in these cell types.

## Results

### Members of the IFN-III and IFN-I families are over-represented in the lower airways of COVID-19 patients

To determine whether IFNs are produced during COVID-19, we studied 152 nasopharyngeal swabs from SARS-CoV-2 positive subjects, including 29 patients with a known clinical follow-up (**Table 1**). In particular, 18 cases required hospitalization, 2 patients needed intensive care unit (ICU) admission, whereas 9 mild cases only required home quarantine. Twenty SARS-CoV-2 negative swabs were included as controls. In the study we also included BALF samples coming from 21 severe hospitalized patients, including 17 ICU-admitted subjects (**Table 1**). Sex and age were distributed among the three groups as reported in **Table 1 and Supplementary Figure 1A-C**.

**TABLE 1.**
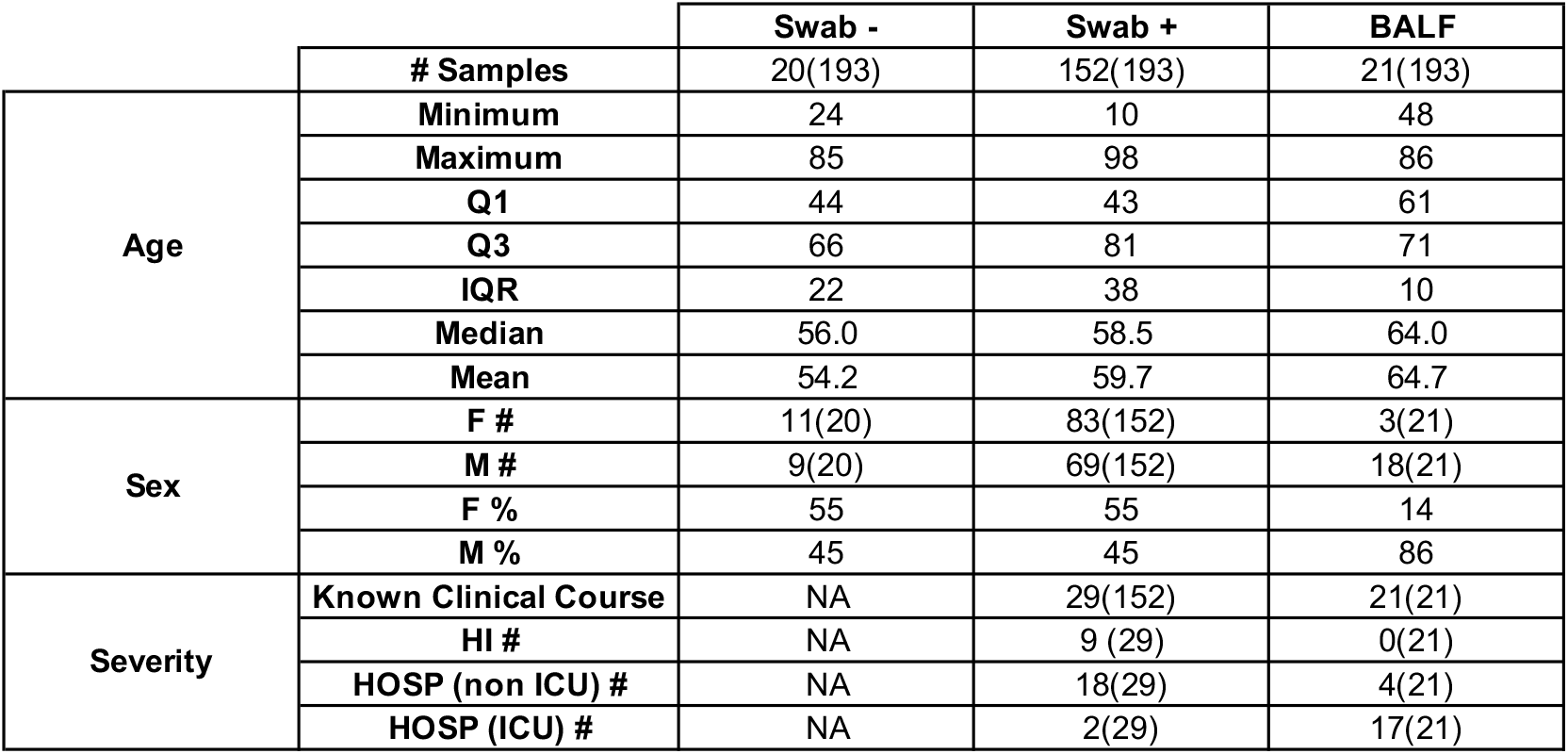
Patient information for Swab −, Swab + and BALF samples for gene expression. Age, sex and severity characteristics of patient cohorts analyzed in **Figure 1**. Nasopharyngeal swabs from SARS-CoV-2-negative (Swab −) and -positive (Swab +) subjects and BALF from SARS-CoV-2-positive patients (BALF) were analyzed. Q1=quartile 1, Q3=quartile 3, IQR=interquartile range, F=female, M=male, HI=home-isolated, HOSP (non ICU)=hospitalized not admitted in the intensive care unit (ICU), HOSP (ICU)=hospitalized and admitted to the ICU, #=number of samples, %=percentage of samples.

Gene expression for the 4 members of the IFN-III family, as well as for selected members of the IFN-I family, was analyzed by quantitative real-time (RT)-PCR in the upper (swabs) and lower (BALF) airways. Transcripts of IFN-III members (with the exception of IFN-λ1) and IFN-I were significantly upregulated in the lower airways compared to the upper airways of patients who tested positive or negative for SARS-CoV-2 (**Figure 1A-F**). Comparison of the production of IFNs in the upper airways of individuals who were SARS-CoV-2 positive versus negative revealed that several IFN members were also significantly upregulated in COVID-19 patients (**Figure 1A-F**). In contrast, no difference was observed in expression of the mRNA for the pro-inflammatory cytokines IL-1β and IL-6 between the three groups (**Figure 1G-H**).

**FIGURE 1.**
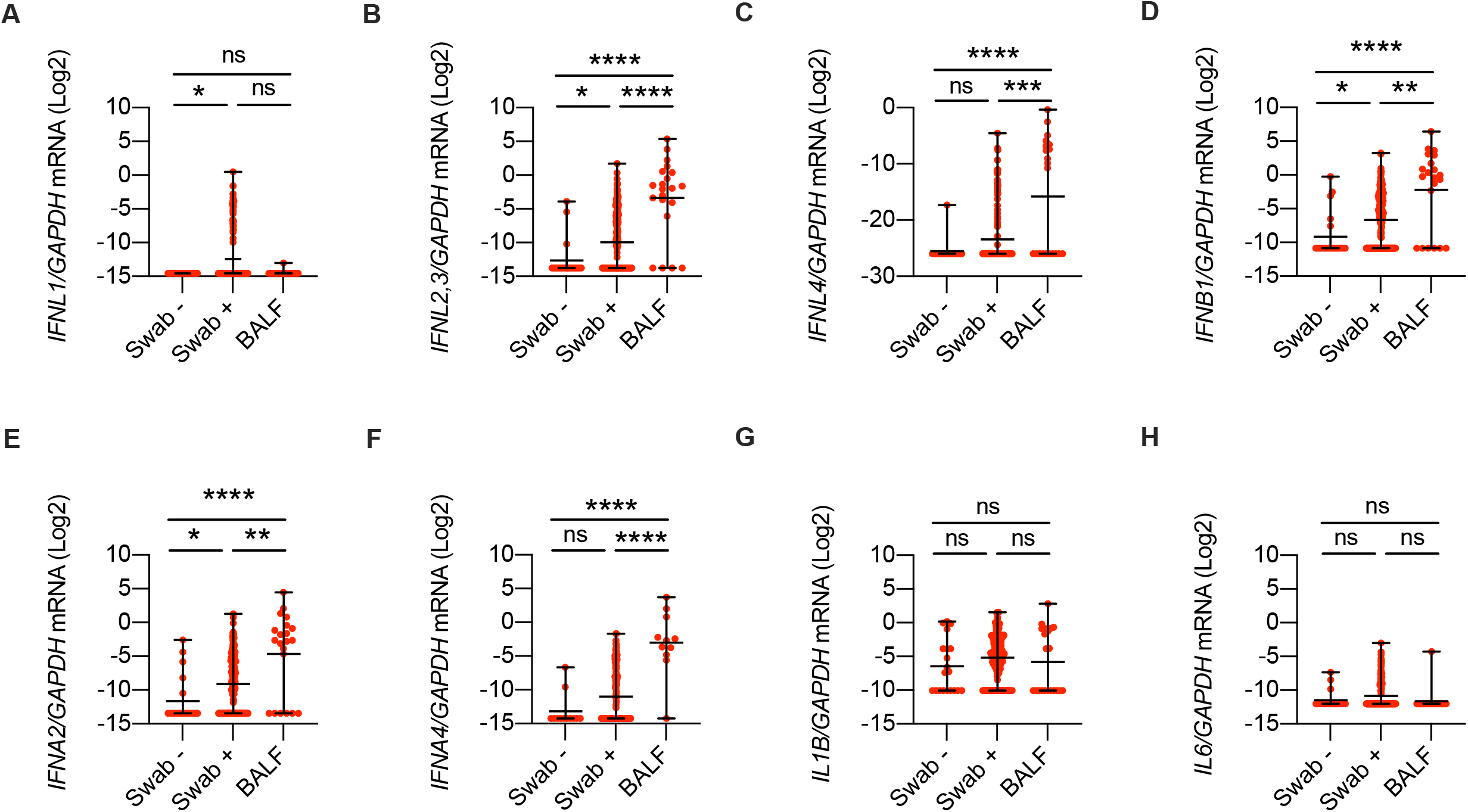
Members of the IFN-III and IFN-I families are over-represented in the lower airways of COVID-19 patients. (**A-H**) *IFNL1* (**A***), IFNL2,3* (**B***), IFNL4* (**C**), *IFNB1* (**D**), *IFNA2* (**E**), *IFNA4* (**F**), *IL1B* (**G**), and *IL6* (**H**) mRNA expression was evaluated in nasopharyngeal swabs from SARS-CoV-2-negative (Swab −) and -positive (Swab +) subjects and in the BALF from SARS-CoV-2-positive patients (BALF). Expression is plotted as log2 (*gene*/*GAPDH* mRNA + 0.5 x gene-specific minimum). Each dot represents a patient. Median with range is depicted. Statistics: (**A-H**) Kruskal-Wallis test with Dunn’s post-hoc test: ns, not significant (*P*>0.05); \**P*<0.05, \*\**P*<0.01, \*\*\**P*<0.001, and \*\*\*\**P*<0.0001.

We observed that gene expression levels of all tested cytokines had a strong bimodal distribution because a fraction of samples had undetectable levels of transcript. We, thus, transformed the gene expression data in categorical variables (expressed or undetected) and evaluated the likelihood of each transcript to be expressed in the BALF as compared to nasopharyngeal swabs (**Supplementary Figure 1D-K**). We found that IFN-λ2,3, IFN-λ4 and IFN-α4 were expressed in a significantly larger proportion of BALF samples compared to SARS-CoV-2 positive swabs (**Supplementary Figure 1E, F, I**). Conversely, transcript of the pro-inflammatory cytokines IL-1β and IL-6 was detected in a significantly larger proportion of nasopharyngeal swabs samples compared to the BALF (**Supplementary Figure 1J-K**). These data demonstrate that selected members of both IFN-III and IFN-I families are over-represented in the lower airways of severe cases of COVID-19, compared to the upper airways of individuals who were infected less severely, or not at all, with SARS-CoV-2.

### A unique IFN signature characterizes the lower airways of COVID-19 patients compared to patients with other ARDS or non-infectious lung pathologies

**Figure 1** shows that the lower airways of severe COVID-19 patients are characterized by the expression of transcripts of several members of the IFN-III and IFN-I families. However, whether the relative distribution of the IFN members, as measured by qPCR, correlates with their protein levels remains unknown. We thus used a multiplex approach to gain insight into the protein levels of IFNs and other inflammatory cytokines in the BALF of subjects infected with COVID-19. We selected an additional 30 patients hospitalized with severe COVID-19 requiring intubation and BALF sampling. The protein signature of these samples was compared to the BALFs of patients with ARDS not driven by SARS-CoV-2 (9, 5 of which were diagnosed with H1N1 influenza A virus infection), and 30 patients with non-infectious lung involvement including fibrosis (10), sarcoidosis (10) or lung transplant (10) (going forward, these will be referred to as “controls”). **Table 2** lists the details of this second cohort of patients. The levels of IFN-III an IFN-I measured in patients with COVID-19 were elevated (**Figure 2A-D**), in agreement with results of the transcriptional analyses (**Figure 1**). IFN-λ2,3 and IFN-α2 were significantly upregulated in COVID-19 patients relative to controls, as well as when compared to patients with ARDS of different etiologies, while the production of IFNλ-1 and IFN-β was significantly increased in COVID-19 patients compared to controls, but not to other ARDS patients (**Figure 2A-D**). Overall, IFN-III were more represented than IFN-I (**Supplementary Figure 2A**). Finally, our results show no correlation between age and IFN levels, nor between age and pro-inflammatory cytokines (**Supplementary Figure 2B-H**).

**FIGURE 2.**
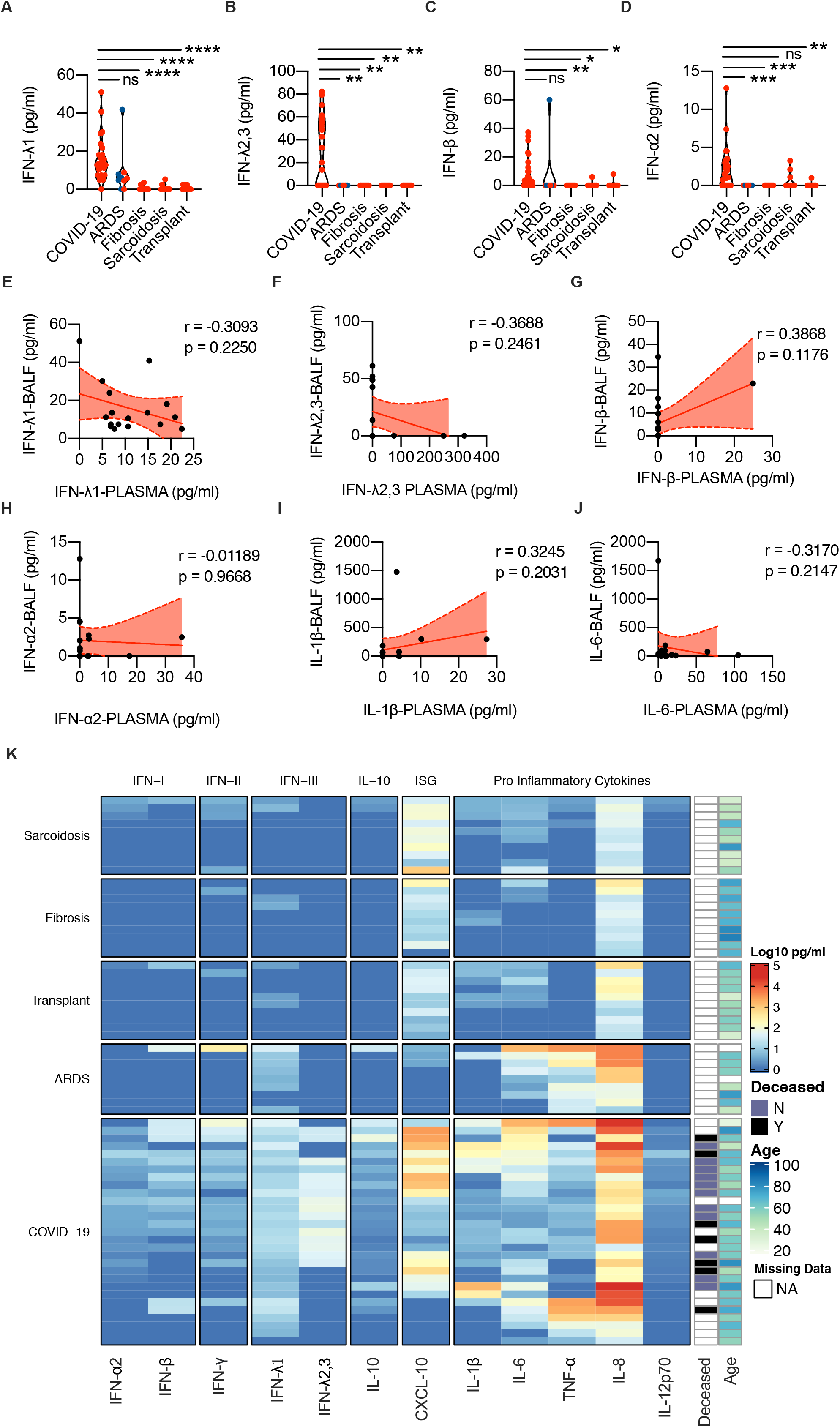
A unique IFN signature characterizes the lower airways of COVID-19 patients compared to patients with other ARDS or non-infectious lung pathologies. (**A-D**) IFN-λ1 (**A**), IFN-λ2,3 (**B**), IFN-β (**C**), IFN-α2 (**D**) protein levels were measured in the BALF of COVID-19, ARDS, Fibrosis, Sarcoidosis, and Transplant patients. Each dot represents a patient. Samples from ARDS patients diagnosed with H1N1 influenza A virus infection are color-coded in blue. Violin plots are depicted. (**E-J**) IFN-λ1 (**E**), IFN-λ2,3 (**F**), IFN-β (**G**), IFN-α2 (**H**), IL-1β (**I**), and IL-6 (**J**) protein levels in the BALF are plotted against protein levels in the plasma of the same patient. Each dot represents a patient. Linear regression lines (continuous line) and 95% confidence interval (dashed line and shaded area) are depicted in red. Spearman correlation coefficients (r) and p-value (p) are indicated. (**K**) Heatmap comparison of IFN-α2, IFN-β, IFN-γ, IFN-λ1, IFN-λ2,3, IL-10, CXCL-10, IL-1β, IL-6, TNF-α, IL-8, IL12p70 protein levels in the BALF of COVID-19, ARDS, Transplant, Fibrosis and Sarcoidosis patients. The color is proportional to the Log10 transformed concentration (pg/ml) of each cytokine. Rows in each group represent different patients. Statistics: (**A-D**) Kruskal-Wallis test with Dunn’s post-hoc test: ns, not significant (*P*>0.05); \**P*<0.05, \*\**P*<0.01, \*\*\**P*<0.001, and \*\*\*\**P*<0.0001. NA: not available.

**TABLE 2.**
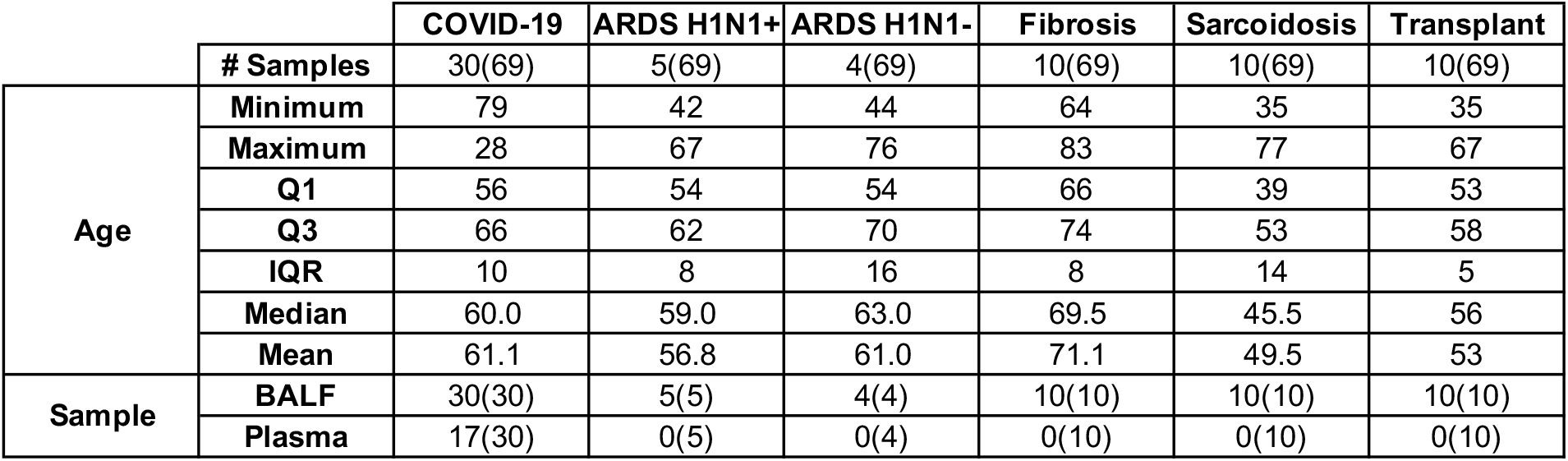
Patient information for BALF and plasma samples used for cytokine quantification. Age and type of collected sample of patient cohorts analyzed in **Figure 2**. BALF and plasma from patients with COVID-19 was analyzed. BALF from patients suffering from: non-COVID-19 ARDS (divided in H1N1 Influenza A virus positive or not), fibrosis, sarcoidosis, and that received lung transplant was analyzed. Q1=quartile 1, Q3=quartile 3, IQR=interquartile range.

Given the recent focus on measuring IFNs or ISGs either in the peripheral blood of COVID-19 patients, or on their local production the lungs (*1, 2, 7, 15–23*), we compared levels of specific members of IFN-III and IFN-I in the BALF and in the plasma of a subset of COVID-19 patients (**Table 2**). We found no correlation between these levels for any of the IFN members studied (**Figure 2E-H**). As with IFN-I and IFN-III, levels of IFN-II and pro-inflammatory cytokines also did not show any correlation between levels found in the BALF and the plasma (**Figure 2I, J, Supplementary Figure 2I**).

We then compared the production in the BALF not only of IFN-III and IFN-I proteins, but also of IFN-II and other proteins associated with the inflammatory process (**Figure 2K** and **Table 3**); we found that COVID-19 patients are characterized by a unique signature of IFNs (which encompasses all three IFN families) and IL-10 production. Many proinflammatory cytokines whose gene induction depends on NF-KB activation are also upregulated in patients who have ARDS that is not driven by SARS-CoV-2; most of these patients also express IFN-λ1, but not other members of the IFN families.

**TABLE 3.**
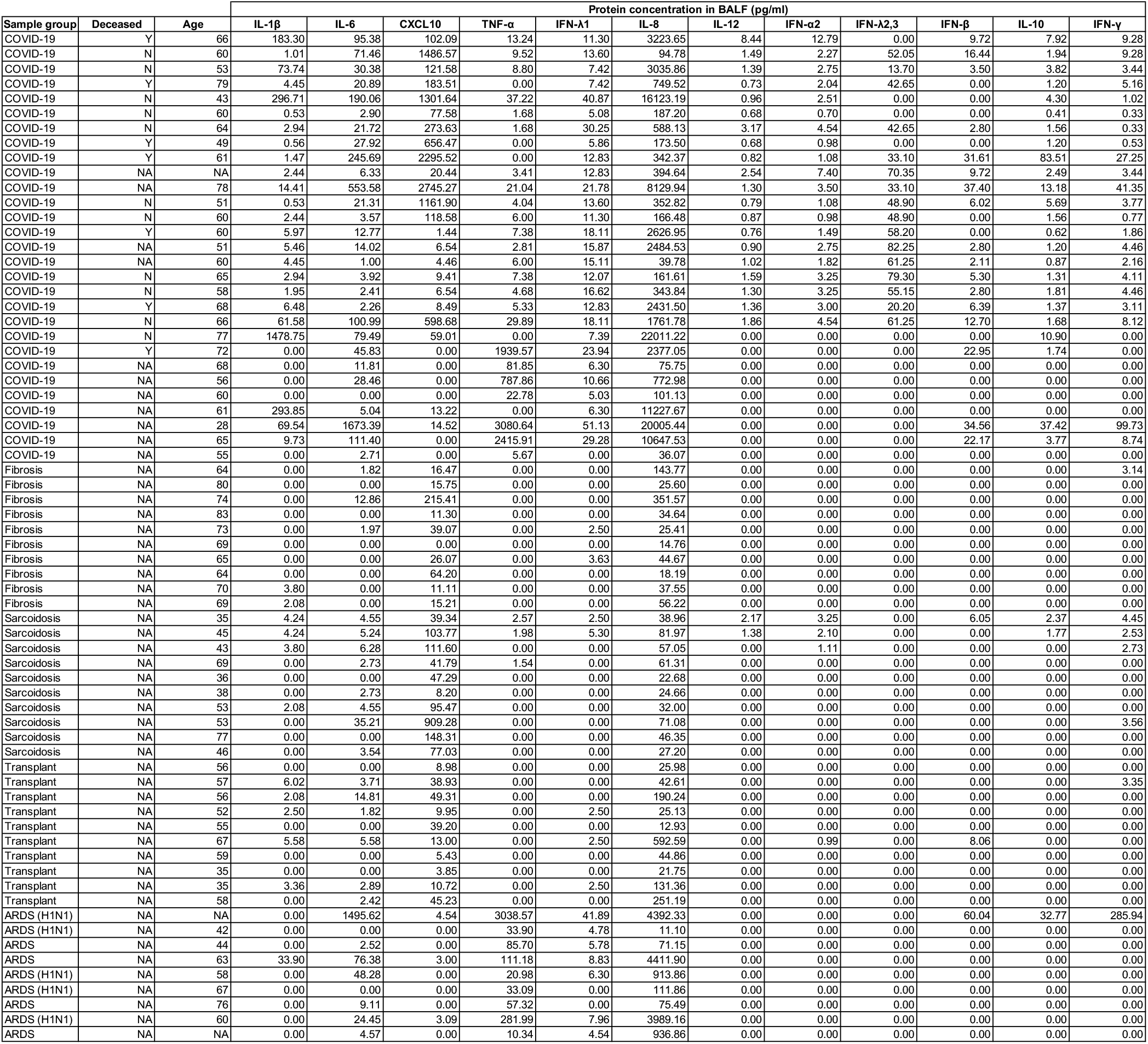
Protein levels in BALF of COVID-19 patients, ARDS patients with non COVID-19 etiology, and patients suffering from non-infectious lung pathologies. Characteristics and protein production levels (pg/ml) of patients analyzed in **Figure 2**. BALF from patients suffering from: non-COVID-19 ARDS (divided in H1N1 Influenza A virus positive or not), fibrosis, sarcoidosis, and that received lung transplant was analyzed. Y=yes, N=no, NA=not available.

Overall, these data demonstrate that COVID-19 patients are characterized by a unique IFN signature in the lower airways relative to patients with ARDS of different etiology, and that IFN-III are the most abundant of all the IFNs in COVID-19 patients.

### High viral loads drive the efficient production of IFN-III, and to a lesser extent of IFN-I, in an age-dependent manner in the upper airways of COVID-19 patients

Our data above (**Figure 1** and **Figure 2**) document the efficient production of IFNs in the lower airways of patients with severe COVID-19. IFNs in general, and IFN-III in particular, are also known to play protective roles in the upper airways during viral infections (*25–27*). Because approximately 15% of severe COVID-19 cases display defects in IFN responses (*20, 21*), we hypothesized that the increased production of IFNs in the lower airways contributes to the cytokine storm, and facilitates detrimental responses (*23, 24*); also that IFN production in the upper airways serves a protective function to prevent the spread of virus into the lower airways and severe disease.

To test this hypothesis, we analyzed gene expression in nasopharyngeal swabs derived from our cohort of 152 SARS-CoV-2 positive patients (Swab+) (**Table 1**) to determine how the pattern of expression of IFNs correlates with viral load, patients’ age, and disease severity. Initially, we examined the distribution of IFN-III and IFN-I levels relative to the viral load, inferred by the mean cycle threshold (Ct) of RT-PCR amplification of the ORF1a/b and E gene regions of SARS-CoV-2 (see Methods for detailed description of viral load definition). We found that expression of several IFNs was correlated with viral levels (**Figure 3A-F**). This is in agreement with another study that reported on ISG levels as a function of viral load in nasopharyngeal swabs (*28*). Of the IFN-III family members, IFN-λ1 and IFN-λ2,3 were positively correlated with viral load. Amongst IFN-I, IFN-β and IFN-α4 also showed a positive correlation with the viral load while IFN-α2 did not show a significant correlation. Transcript levels of the proinflammatory cytokines *IL1B* and *IL6* were also positively correlated to the viral load (**Figure 3G, H**).

**FIGURE 3.**
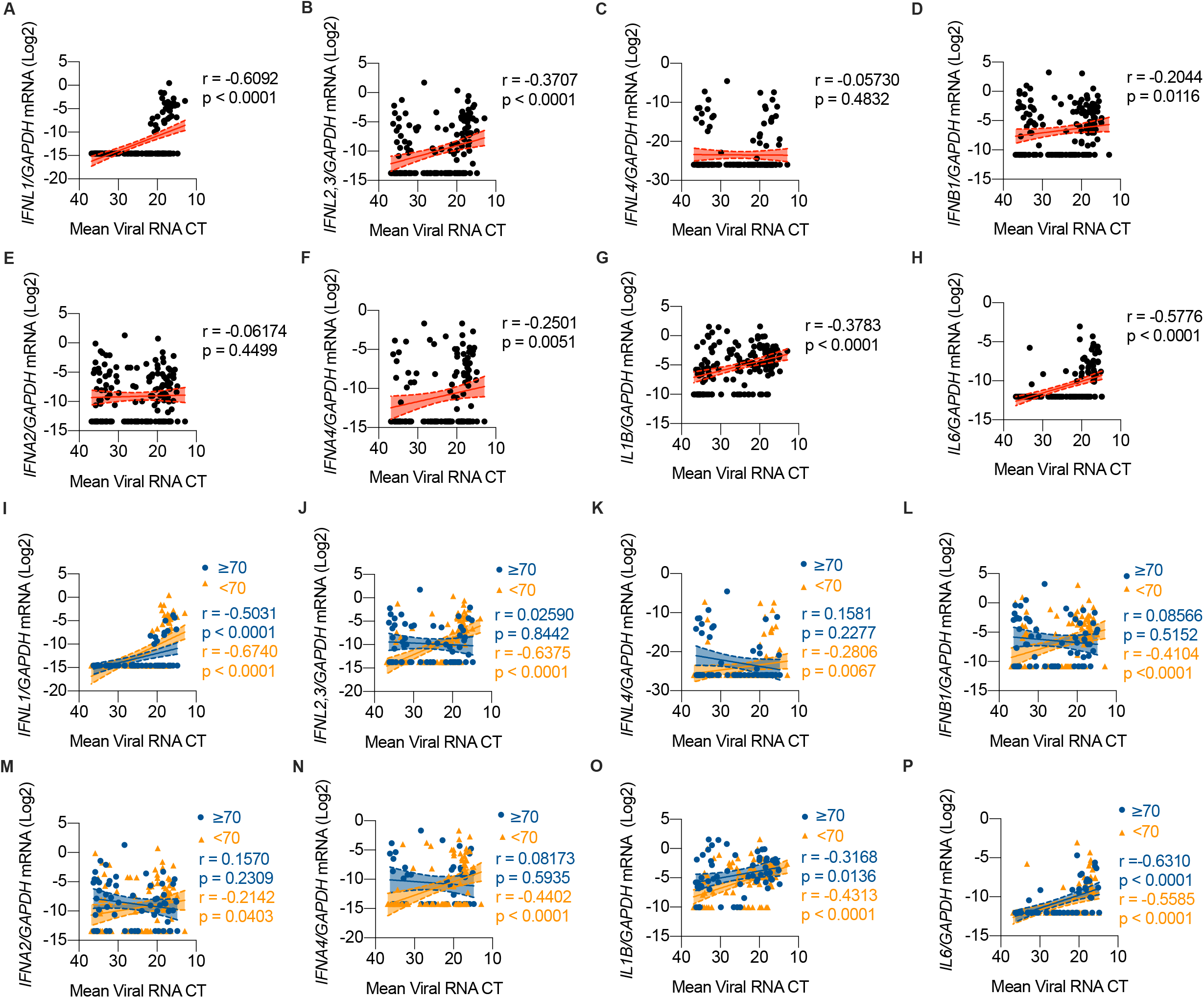
High viral loads drive the efficient production of IFN-III, and to a lesser extent of IFN-I, in an age-dependent manner in the upper airways of COVID-19 patients. (**A-H**) *IFNL1* (A*), IFNL2,3* (B*), IFNL4* (**C**), *IFNB1* (**D**), *IFNA2* (**E**), *IFNA4* (**F**), *IL1B* (**G**), and *IL6* (**H**) mRNA expression is plotted against mean viral RNA CT in SARS-CoV-2^+^ swabs. (**A-H**) Expression is plotted as log2 (*gene*/*GAPDH* mRNA *+* 0.5 x gene-specific minimum). Each dot represents a patient. Linear regression lines (continuous line) and 95% confidence interval (dashed line and shaded area) are depicted in red. Spearman correlation coefficients (r) and p-value (p) are indicated. (**I-P**) *IFNL1* (I*), IFNL2,3* (J*), IFNL4* (**K**), *IFNB1* (**L**), *IFNA2* (**M**), *IFNA4* (**N**), *IL1B* (**O**), and *IL6* (**P**) mRNA expression is plotted against mean viral RNA CT in swabs from SARS-CoV-2^+^ patients over 70-year-old (≥ 70, blue dots and lines) and below 70-year-old (< 70, orange dots and lines). (**I-P**) Expression is plotted as log2 (*gene*/*GAPDH* mRNA *+* 0.5 x gene-specific minimum). Each dot represents a patient. Linear regression (continuous lines) and 95% confidence interval (dashed line and shaded area) are depicted. Spearman correlation coefficients (r) and p-value (p) are indicated in blue and in orange for ≥70 and <70 year-old patients respectively.

We next evaluated how gene expression relates to the age of patients. Several studies have shown that the severity and lethality of COVID-19 increase with age, with a sharp upturn starting in the eighth decade (*29, 30*). In particular, elderly patients display prolonged and dysregulated production of IFN-I, II and III in the plasma (*1*). We thus divided our cohort of patients into those that were greater than or less than 70 years of age (indicated as “≥70”, and “<70”, respectively). Our analyses confirmed that IFN-λ1 expression is tightly associated with the viral load and is independent of patients’ age (**Figure 3I**). In contrast, expression of IFN-λ2,3 and IFN-λ4 maintained the association with the viral load only for the < 70 cohort (**Figure 3J, K**). Also, IFN-I showed an association with the viral load in the <70, but not ≥70, patient cohort (**Figure 3L-N**), while both pro-inflammatory cytokines maintained their association with the viral load independent of age (**Figure 3O, P**).

Next, we divided our patient cohort into terciles based on the viral load (**Table 4**), and analyzed gene expression. Results confirmed that at the highest viral load, IFN-λ2,3 and IFN-β are significantly upregulated in the <70, compared to the ≥70, cohort (**Supplementary Figure 3A-F**). Notably, while no difference was observed between age groups for *IL-6*, elder patients (≥70) showed significant increase in *IL1B* levels compared to younger ones at low viral loads (**Supplementary Figure 3G, H**). These findings confirm, also at the local level, the complex picture of the inflammatory signatures that have been reported for blood of COVID-19 patients (*1*) and further support the importance of IL-1β in driving detrimental immune responses during lung viral infections (*31*).

**TABLE 4.**
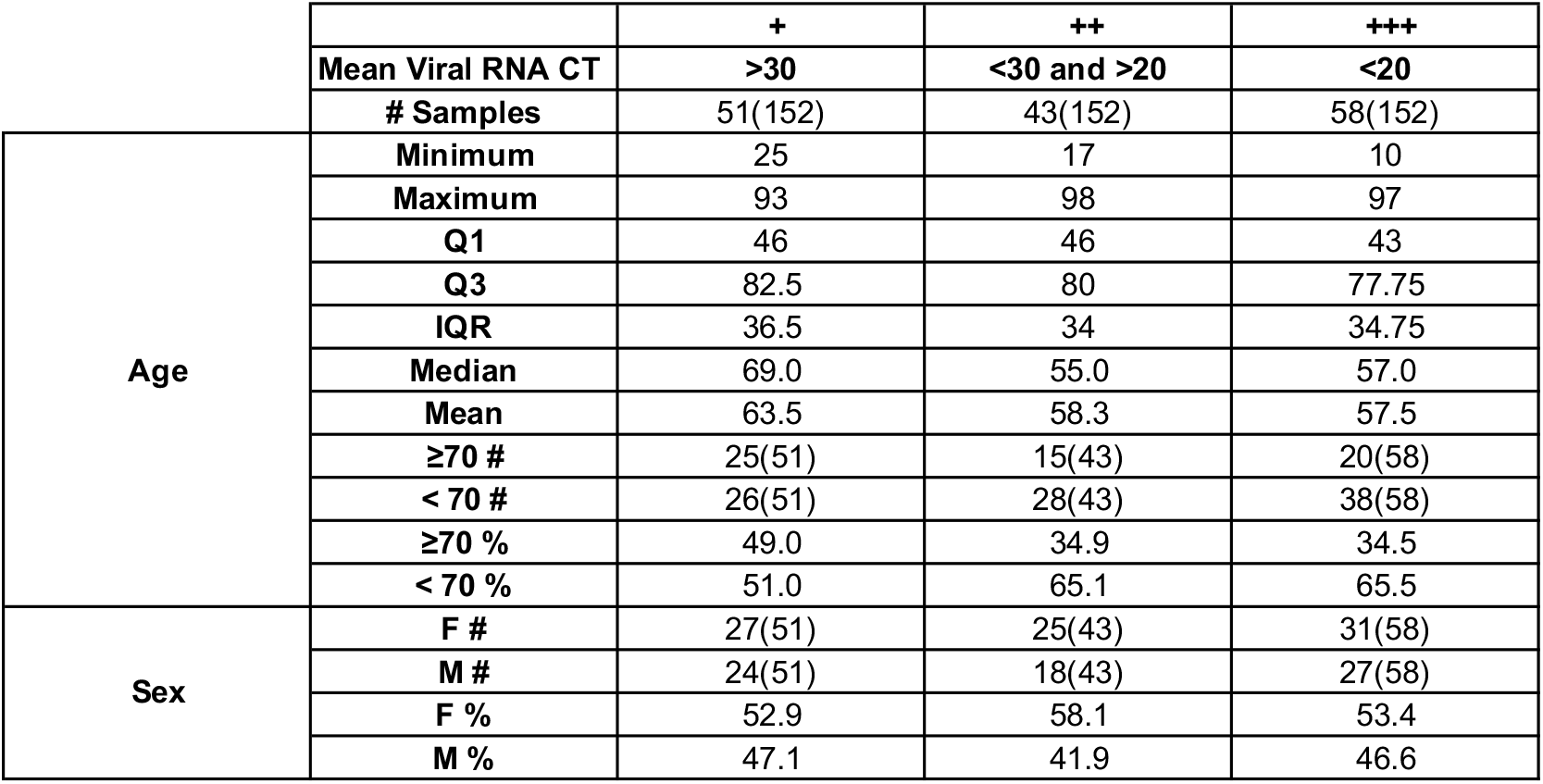
Patient information for +, ++, +++ swab samples used for gene expression. Age and sex characteristics of patient cohorts analyzed in **Figure 3 and Supplementary Figure 3**. Nasopharyngeal swabs from SARS-CoV-2^+^ patients divided in viral load terciles (“+++”, “++”, “+”) were analyzed. Q1=quartile 1, Q3=quartile 3, IQR=interquartile range, ≥ 70=over or equal to 70 years old, <70=under 70 years old, F=female, M=male, #=number of samples, %=percentage of samples.

|  |  | + | ++ | +++ |
| --- | --- | --- | --- | --- |
|  | Mean Viral RNA CT | >30 | <30 and >20 | <20 |
|  | # Samples | 51(152) | 43(152) | 58(152) |
| Age | Minimum | 25 | 17 | 10 |
|  | Maximum | 93 | 98 | 97 |
|  | Q1 | 46 | 46 | 43 |
|  | Q3 | 82.5 | 80 | 77.75 |
|  | IQR | 36.5 | 34 | 34.75 |
|  | Median | 69.0 | 55.0 | 57.0 |
|  | Mean | 63.5 | 58.3 | 57.5 |
|  | ≥70 # | 25(51) | 15(43) | 20(58) |
|  | < 70 # | 26(51) | 28(43) | 38(58) |
|  | ≥70 % | 49.0 | 34.9 | 34.5 |
|  | < 70 % | 51.0 | 65.1 | 65.5 |
| Sex | F # | 27(51) | 25(43) | 31(58) |
|  | M # | 24(51) | 18(43) | 27(58) |
|  | F % | 52.9 | 58.1 | 53.4 |
|  | M % | 47.1 | 41.9 | 46.6 |

To account for the bimodal distribution of cytokine gene expression data we transformed it in discrete variables (expressed or undetected). Using logistic regression, we estimated the odds ratio for genes to be expressed in medium (“++”) and high viral (“+++”) load samples compared to low viral load (“+”) and tested if there was an interaction between viral load and age groups on gene expression. We found that response patterns to viral load were significantly different between elder (≥70) and younger (<70) patients for IFN-λ2,3 (P value for interaction<0.001) and IFN-α4 (P value for interaction=0.03). This analysis also showed that only younger patients have a dose-response relationship between IFN gene expression and viral load (**Supplementary Figure 3I-M and Table 5**), further confirming a dysregulated production of IFNs in the elderly (≥70) group, not only at the systemic (*1*), but also at the local level. In contrast to IFNs, no difference in the dose-response relationship between IL-1β and IL-6 expression and viral load was observed between age groups **(Supplementary Figure 3N, O, G, H and Table 5**).

**TABLE 5.** Odds ratio of expressing/not each gene across viral load terciles and age groups in SARS-CoV-2 + swabs. Odds ratio of expressing *IFNL2,3*, *IFNL4*, *IFNB1*, *IFNA2*, *IFNA4*, *IL1B*, and *IL6* mRNA in “+++” and “++” with respect to “+” SARS-CoV-2^+^ swabs in ≥ 70 and < 70 patients was calculated. Odds ratio column indicates the odds ratio and associated 95% confidence interval in brackets. P value column indicates the associated P value for each cohort of patients. Interaction between viral load terciles and age groups (≥70 years vs <70 years) was tested and P values for interaction are indicated. NE=not estimable.

| Outcome/gene | Viral load, mean CT | Age <70 |  | Age ≥70 |  | P value for interaction |
| --- | --- | --- | --- | --- | --- | --- |
|  |  | Odds ratio | P value | Odds ratio | P value |  |
| *IFNA2* | + | Reference(1.0) |  | Reference(1.0) |  | 0.38 |
|  | ++ | 0.7(0.2,2.2) | 0.56 | 0.5(0.1,1.9) | 0.29 |  |
|  | +++ | 3.2(1.1,9.2) | 0.03 | 0.9(0.2,3.6) | 0.88 |  |
| *IFNA4* | + | Reference(1.0) |  | Reference(1.0) |  | 0.03 |
|  | ++ | 1.2(0.2,6.1) | 0.83 | 0.6(0.1,2.8) | 0.49 |  |
|  | +++ | 10.8(2.4,48.8) | 0.001 | 0.8(0.2,3.0) | 0.69 |  |
| *IFNB1* | + | Reference(1.0) |  | Reference(1.0) |  | 0.15 |
|  | ++ | 0.4(0.1,1.4) | 0.16 | 0.4(0.1,1.7) | 0.23 |  |
|  | +++ | 8.8(2.6,30.1) | <0.001 | 1.6(0.4,6.8) | 0.49 |  |
| *IFNL1* | + | Reference(1.0) |  | Reference(1.0) |  |  |
|  | ++ | NE |  | NE |  |  |
|  | +++ | NE |  | NE |  |  |
| *IFNL2,3* | + | Reference(1.0) |  | Reference(1.0) |  | <0.001 |
|  | ++ | 0.7(0.2,2.9) | 0.60 | 0.8(0.2,2.8) | 0.68 |  |
|  | +++ | 22.1(6.0,82.0) | <0.001 | 1.0(0.4,3.5) | 0.96 |  |
| *IFNL4* | + | Reference(1.0) |  | Reference(1.0) |  | 0.13 |
|  | ++ | NE |  | 0.6(0.2,2.6) | 0.54 |  |
|  | +++ | 4.2(1.0,17.6) | 0.049 | 0.6(0.2,2.1) | 0.39 |  |
| *IL1B* | + | Reference(1.0) |  | Reference(1.0) |  | 0.08 |
|  | ++ | 6.1(1.7,21.2) | 0.005 | 0.6(0.1,3.2) | 0.61 |  |
|  | +++ | 11.4(3.1,42.1) | <0.001 | NE |  |  |
| *IL6* | + | Reference(1.0) |  | Reference(1.0) |  | 0.82 |
|  | ++ | 2.9(0.3,30.1) | 0.36 | 5.9(0.5,63.8) | 0.14 |  |
|  | +++ | 28.1(3.4,233.1) | 0.002 | 80.0(7.9,805.8) | <0.001 |  |

### Mild COVID-19 is characterized by high levels of IFN-III, not IFN-I, in response to high viral loads in the upper airways

The data in **Figure 3** suggest that younger patients, associated with lower risk to develop severe COVID-19, present higher levels of IFNs when the viral load is high. To explore a possible link between IFN production and disease severity, we repeated our analyses in our subset of patients with a known clinical follow-up. Disease severity was assessed as follows: hospitalized patients (HOSP) that were further grouped between those admitted to the ICU (HOSP ICU) and not (HOSP non ICU), and patients that have been discharged from the emergency room without being hospitalized that were identified as home-isolated (HI) (**Table 6**). When IFN-III levels were plotted against the viral load, only the less severe, home isolated patients showed a positive correlation with IFN-λ2,3 expression, while the viral load correlated with IFN-λ1 in both groups (**Figure 4A-C**); this correlation was lost for IFN-I (**Figure 4D-F**). With regard to pro-inflammatory cytokines, the positive correlation between *IL6* levels and viral load was maintained only for hospitalized patients, while we could not observe positive correlation between viral load and *IL1B* expression in either of the patient groups (**Figure 4G, H**).

**FIGURE 4.**
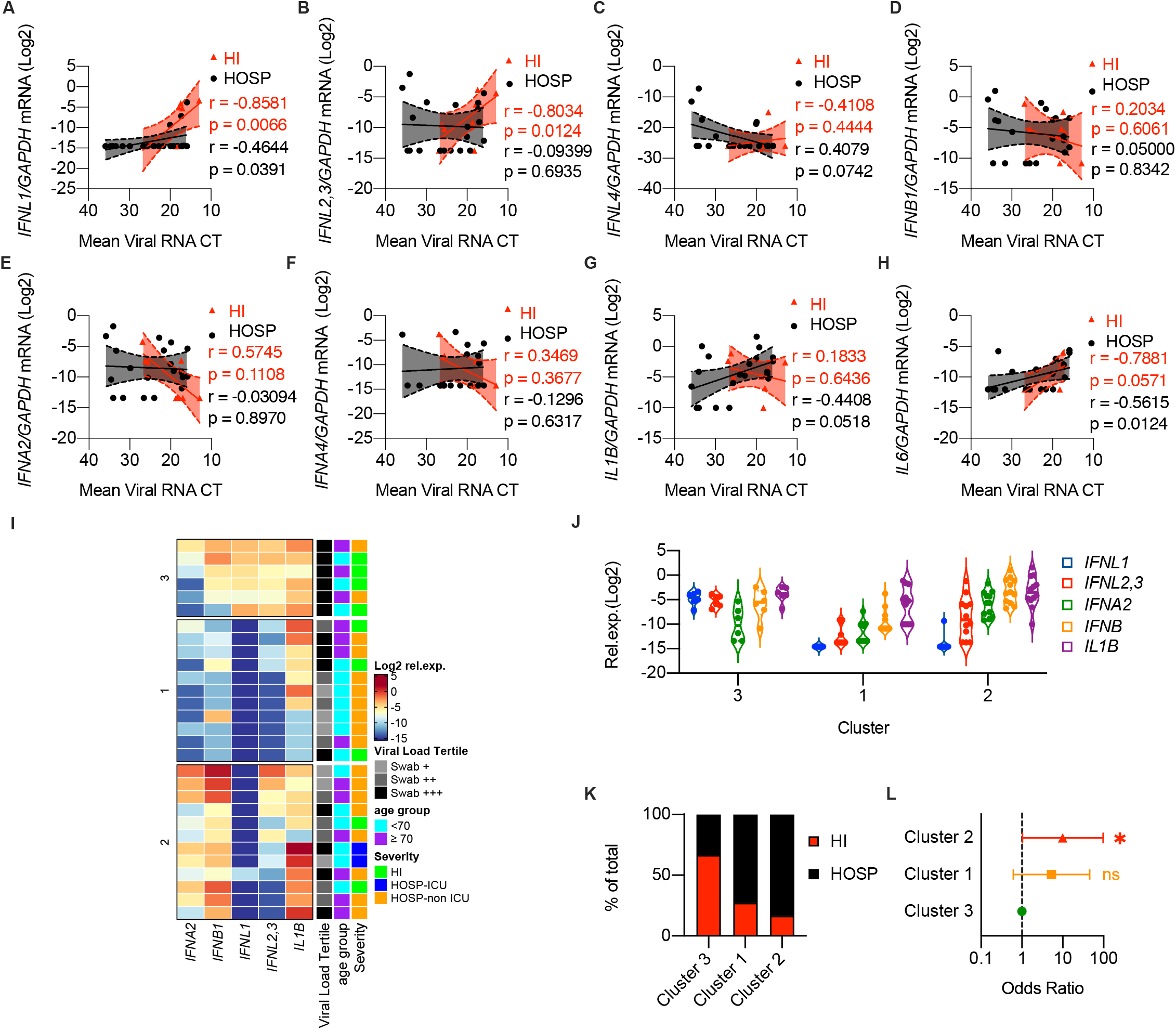
Mild COVID-19 is characterized by high levels of IFN-III, not IFN-I, in response to high viral loads in the upper airways. (**A-L**) Swabs from a cohort of SARS-CoV-2^+^ patients with known disease severity including ICU inpatients and hospitalized patients (HOSP, black dots and lines) and home-isolated patients (HI, red dots and lines) were analyzed. (**A-H**) *IFNL1* (**A***), IFNL2,3* (**B***), IFNL4* (**C**), *IFNB1* (**D**), *IFNA2* (**E**), *IFNA4* (**F**), *IL1B* (**G**), and *IL6* (**H**) mRNA expression is plotted against mean viral RNA CT. Expression is plotted as log2 (*gene*/*GAPDH* mRNA *+* 0.5 x gene-specific minimum). Each dot represents a patient. Linear regression lines (continuous line) and 95% confidence interval (dashed line and shaded area) are depicted. Spearman correlation coefficients (r) and p-value (p) are indicated in black and in red for HOSP and HI patients respectively. (**I**) K-means clustering based on the expression of *IFNA2, IFNB1 IFNL1, IFNL2,3*, *IL1B* was used to determine clusters 1-3 (Cluster 1 n=15, Cluster 2 n=8, Cluster 3 n=6). The color indicates the relative gene expression. Viral load tercile, age group and severity are annotated. Viral load terciles (“+++”, “++”, “+”) are defined by mean viral RNA CT (<20, >20 and <30, > 30). Age groups are defined as <70 or ≥70-year old patients. Severity groups are defined as follows: HI=home isolated, HOSP (non ICU)=Hospitalized patients that did not require ICU admission, HOSP (ICU)=Hospitalized patients admitted to the ICU. (**J**) *IFNL1, IFNL2,3*, *IFNA2, IFNB1, IL1B* mRNA expression within clusters identified in Figure 4I. Expression is plotted as log2 (*gene*/*GAPDH* mRNA *+* 0.5 x gene-specific minimum). Each dot represents a patient. Violin plots are depicted. (**K**) Percentage of patients with the indicated disease severity within clusters identified in Figure 4I. (**L**) Odds ratio of patients in Cluster 2 and Cluster 1 being hospitalized relative to patients in Cluster 3 (Clusters identified in Figure 4I). Symbols represent the odds ratio. Error bars represent the 95% confidence interval associated to the odds ratio. Statistics: (**L**) Odds ratio: ns, not significant (*P*>0.05); \**P*<0.05, \*\**P*<0.01, \*\*\**P*<0.001.

**TABLE 6.**
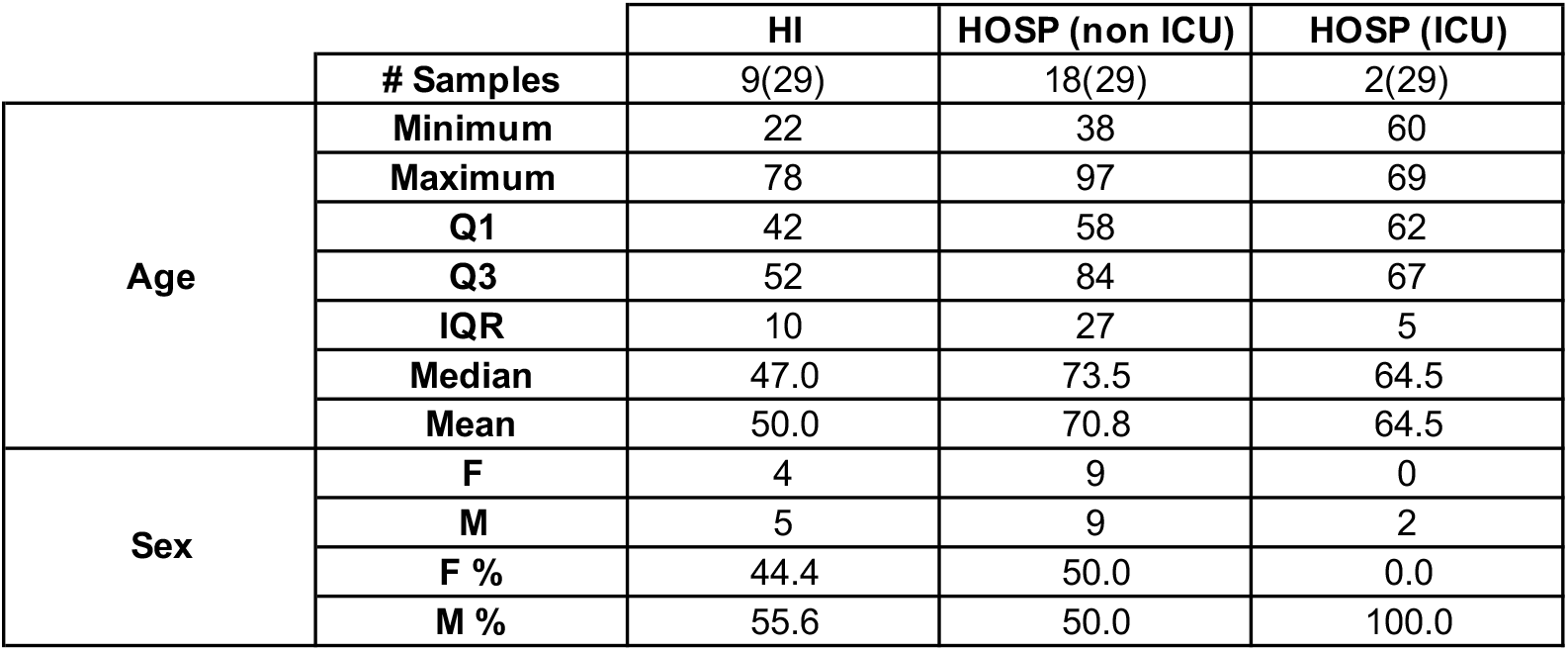
Patient information for HI, HOSP (non ICU), HOSP (ICU) swab samples used for gene expression. Age and sex characteristics of patient cohorts analyzed in **Figure 4**. Nasopharyngeal swabs from SARS-CoV-2 positive patients home-isolated (HI), hospitalized not admitted in the intensive care unit (HOSP (non ICU)) and hospitalized and admitted to the ICU (HOSP (ICU)) were analyzed. Q1=quartile 1, Q3=quartile 3, IQR=interquartile range, F=female, M=male, #=number of samples, %=percentage of samples.

We then divided these samples into 3 clusters via unbiased K-mean clustering, based on their expression of IFN-I, IFN-III and the proinflammatory cytokine IL-1β. Our results reveal that cluster 3, characterized by high expression of IFN-III (IFN-λ1, IFN-λ2,3), and to a lesser extent of IFN-I, was enriched in home-isolated patients with milder disease manifestations and high viral load (**Figure 4I-K**, **Supplementary Figure 4A-C**). In contrast, cluster 2 (low levels of IFN-III and the highest levels of IFN-I) and cluster 1 (low IFN-I and IFN-III expression and high IL-1β expression) were enriched in the most severe patients (**Figure 4I-K**, **Supplementary Figure 4A**). Moreover, patients in cluster 2 were 10 times more likely to have severe illness resulting in hospitalization or ICU admission than patients in cluster 3 (**Figure 4L,** odds ratio 10.1, 95% confidence interval 1.0-97.5, P value <0.05). Overall, these data support the hypothesis that efficient production of IFN-III in the upper airways of COVID-19 patients with high viral load protects against severe COVID-19.

### Bronchial epithelial cells and phagocytes produce specific members of the IFN-III and IFN-I family when activated by different PRRs

A general consensus in the field is that tropism of SARS-CoV-2 favors the infection of ACE2-positive epithelial cells along the respiratory tract (*32–35*), while immune cells respond during COVID-19 to either viral components or to host molecules released during the infection process (*33, 35–37*). Recently, activation of the retinoic acid-inducible gene I (RIG-I)/ melanoma differentiation-associated protein 5 (MDA-5) pathway in epithelial cells has been implicated as a major driver of IFN production in response to SARS-CoV-2 (*38*). Based on our data, we hypothesized that different populations of cells contribute to IFN production during a viral infection by activating specific PRRs. We also found that, while the mRNA for *IFNL1* is absent in the lower airways of COVID-19 patients (**Figure 1A**), protein levels for IFN-λ1 are abundant at the same anatomical site (**Supplementary Figure 2A**). These data suggest that cells that actively produce the mRNA for *IFNL1* are underrepresented in the BALF. However, *IFNL1* is one of the most upregulated genes in the upper airways. Given that epithelial cells are over-represented in swabs from upper epithelial airways (*35*), but are under-represented in the BALF (*39, 40*), we next explored how PRR activation of epithelial cells, or of phagocytes, differentially drives IFN production (**Figure 5A**).

**FIGURE 5.**
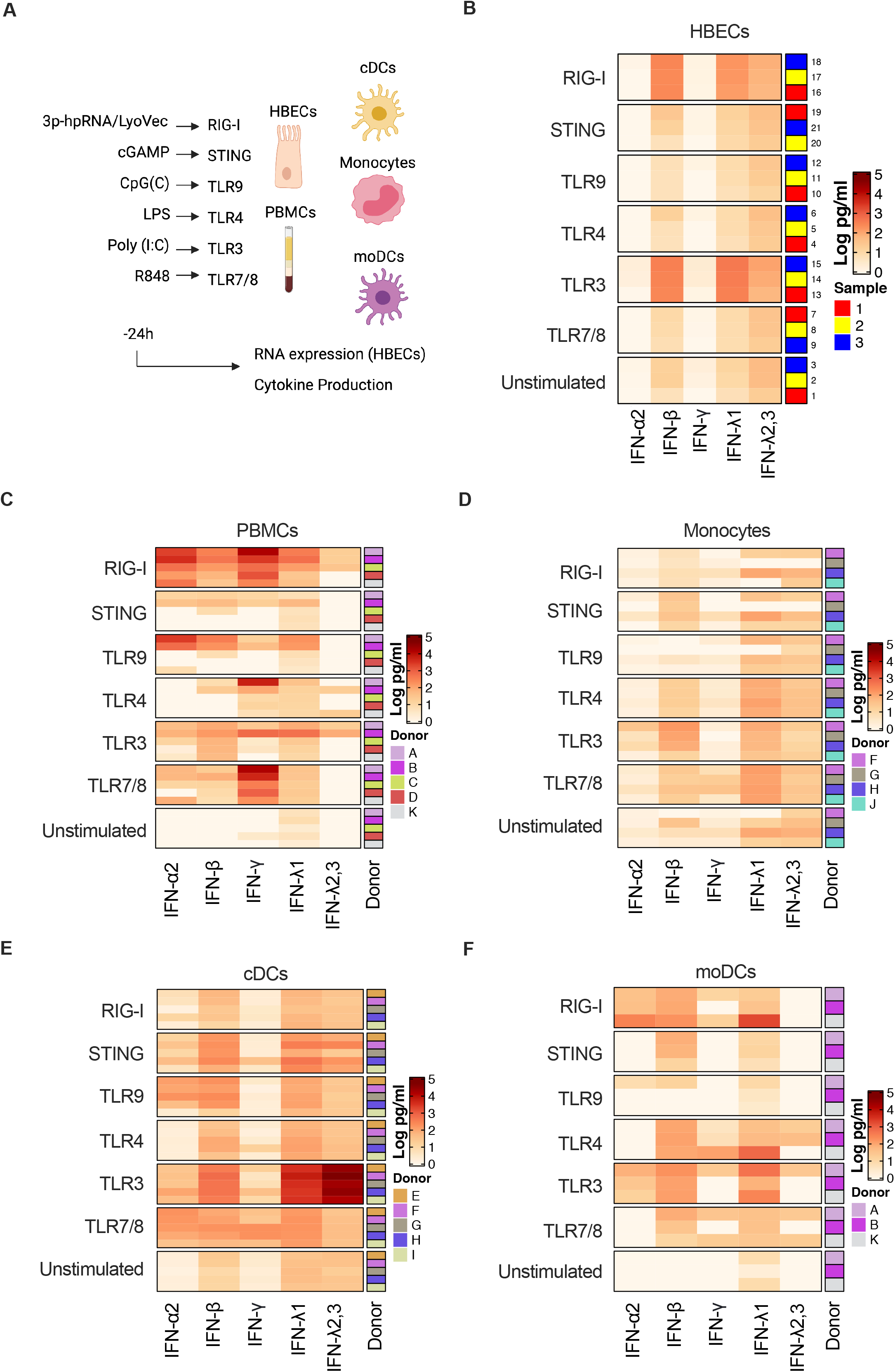
Bronchial epithelial cells and phagocytes produce specific members of the IFN-III and IFN-I family when activated by different PRRs. **(A**) Schematic of experimental setup. HBECs, PBMCs, Monocytes, cDCs and moDCs were treated for 24 hours with 3p-hpRNA/LyoVec, cGAMP, CpG(C), LPS, Poly (I:C), R848 for stimulation of RIG-I, STING, TLR9, TLR4, TLR3, TLR7/8 respectively. Cytokine expression was evaluated on RNA extracted from cell lysates and cytokine production was evaluated in supernatants (created with BioRender). (**B-F**) Heatmap representation of IFN-α2, IFN-β, IFN-γ, IFN-λ1 and IFN-λ2,3 production by HBECs (**B**), PMBCs (**C**), Monocytes (**D**), cDCs (**E**), moDCs (**F**) 24 hours after treatment. The color is proportional to the Log10 transformed concentration (pg/ml) of each cytokine. (**B**) Rows in each group represent a biological replicate. (**C-F**) Rows in each group represent different donors as depicted in the annotation.

Our data confirmed that of the PRRs targeted to activate epithelial cells, the RIG-I pathway, and to a lesser extent TLR3, were the most potent inducers of IFNs, and in particular of IFN-λ1, IFN-λ2,3 and IFN-β, at the mRNA level in epithelial cells (**Supplementary Figure 5A, Table 7A**). Also, epithelial cells showed a preferential production of IFNs, and consequently of ISGs, relative to other pro-inflammatory mediators (**Supplementary Figure 5A, Table 7A**). A comparison of the protein levels for the different IFNs produced by epithelial cells revealed that TLR3 and RIG-I stimulation led to an efficient production of both IFN-III and IFN-I (**Figure 5B, Supplementary Figure 5C, Table 7B**), and to a lesser extent of inflammatory cytokines (**Supplementary Figure 5B, Table 7B**). Notably, epithelial cells are more potent producers of IFN-λ1 compared to IFN-λ2,3 upon stimulation of TLR3, RIG-I and MDA-5 pathways with Poly (I:C), 3p-hpRNA and transfected Poly (I:C) respectively (**Figure 5B, Supplementary Figure 5C**).

**TABLE 7.**
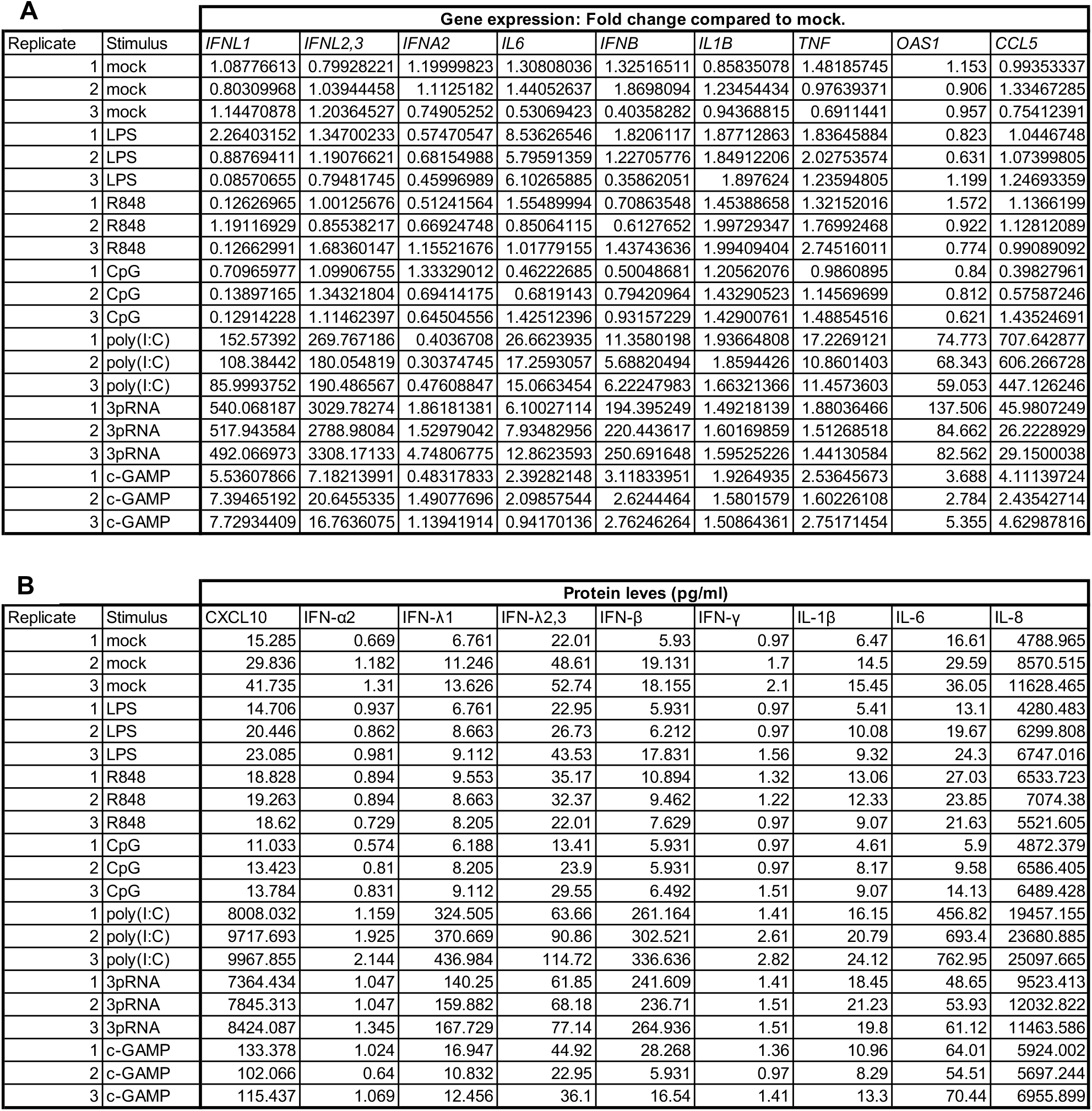
Cytokine gene expression and protein production from human bronchial epitheial cells in response to a panel of PAMPs. Cytokine gene expression levels (expressed as Fold Change compared to mock) (**A**) and cytokine protein levels (pg/ml) (**B**) produced by HBECs stimulated with agonists of PRRs in three biological triplicates as described in **Figure 5**.

**A**
|  |  | Gene expression: Fold change compared to mock. |  |  |  |  |  |  |  |  |
| --- | --- | --- | --- | --- | --- | --- | --- | --- | --- | --- |
| Replicate | Stimulus | *IFNL1* | *IFNL2,3* | *IFNA2* | *IL6* | *IFNB* | *IL1B* | *TNF* | *OAS1* | *CCL5* |
| 1 | mock | 1.08776613 | 0.79928221 | 1.19999823 | 1.30808036 | 1.32516511 | 0.85835078 | 1.48185745 | 1.153 | 0.99353337 |
| 2 | mock | 0.80309968 | 1.03944458 | 1.1125182 | 1.44052637 | 1.8698094 | 1.23454434 | 0.97639371 | 0.906 | 1.33467285 |
| 3 | mock | 1.14470878 | 1.20364527 | 0.74905252 | 0.53069423 | 0.40358282 | 0.94368815 | 0.6911441 | 0.957 | 0.75412391 |
| 1 | LPS | 2.26403152 | 1.34700233 | 0.57470547 | 8.53626546 | 1.8206117 | 1.87712863 | 1.83645884 | 0.823 | 1.0446748 |
| 2 | LPS | 0.88769411 | 1.19076621 | 0.68154988 | 5.79591359 | 1.22705776 | 1.84912206 | 2.02753574 | 0.631 | 1.07399805 |
| 3 | LPS | 0.08570655 | 0.79481745 | 0.45996989 | 6.10265885 | 0.35862051 | 1.897624 | 1.23594805 | 1.199 | 1.24693359 |
| 1 | R848 | 0.12626965 | 1.00125676 | 0.51241564 | 1.55489994 | 0.70863548 | 1.45388658 | 1.32152016 | 1.572 | 1.1366199 |
| 2 | R848 | 1.19116929 | 0.85538217 | 0.66924748 | 0.85064115 | 0.6127652 | 1.99729347 | 1.76992468 | 0.922 | 1.12812089 |
| 3 | R848 | 0.12662991 | 1.68360147 | 1.15521676 | 1.01779155 | 1.43743636 | 1.99409404 | 2.74516011 | 0.774 | 0.99089092 |
| 1 | CpG | 0.70965977 | 1.09906755 | 1.33329012 | 0.46222685 | 0.50048681 | 1.20562076 | 0.9860895 | 0.84 | 0.39827961 |
| 2 | CpG | 0.13897165 | 1.34321804 | 0.69414175 | 0.6819143 | 0.79420964 | 1.43290523 | 1.14569699 | 0.812 | 0.57587246 |
| 3 | CpG | 0.12914228 | 1.11462397 | 0.64504556 | 1.42512396 | 0.93157229 | 1.42900761 | 1.48854516 | 0.621 | 1.43524691 |
| 1 | poly(I:C) | 152.57392 | 269.767186 | 0.4036708 | 26.6623935 | 11.3580198 | 1.93664808 | 17.2269121 | 74.773 | 707.642877 |
| 2 | poly(I:C) | 108.38442 | 180.054819 | 0.30374745 | 17.2593057 | 5.68820494 | 1.8594426 | 10.8601403 | 68.343 | 606.266728 |
| 3 | poly(I:C) | 85.9993752 | 190.486567 | 0.47608847 | 15.0663454 | 6.22247983 | 1.66321366 | 11.4573603 | 59.053 | 447.126246 |
| 1 | 3pRNA | 540.068187 | 3029.78274 | 1.86181381 | 6.10027114 | 194.395249 | 1.49218139 | 1.88036466 | 137.506 | 45.9807249 |
| 2 | 3pRNA | 517.943584 | 2788.98084 | 1.52979042 | 7.93482956 | 220.443617 | 1.60169859 | 1.51268518 | 84.662 | 26.2228929 |
| 3 | 3pRNA | 492.066973 | 3308.17133 | 4.74806775 | 12.8623593 | 250.691648 | 1.59525226 | 1.44130584 | 82.562 | 29.1500038 |
| 1 | c-GAMP | 5.53607866 | 7.18213991 | 0.48317833 | 2.39282148 | 3.11833951 | 1.9264935 | 2.53645673 | 3.688 | 4.11139724 |
| 2 | c-GAMP | 7.39465192 | 20.6455335 | 1.49077696 | 2.09857544 | 2.6244464 | 1.5801579 | 1.60226108 | 2.784 | 2.43542714 |
| 3 | c-GAMP | 7.72934409 | 16.7636075 | 1.13941914 | 0.94170136 | 2.76246264 | 1.50864361 | 2.75171454 | 5.355 | 4.62987816 |

**TABLE 7. Cytokine gene expression and protein production from human bronchial epithelial cells in response to a panel of PAMPs.**
|  |  | Protein levels (pg/ml) |  |  |  |  |  |  |  |  |
| --- | --- | --- | --- | --- | --- | --- | --- | --- | --- | --- |
| Replicate | Stimulus | CXCL10 | IFN-α2 | IFN-λ1 | IFN-λ2,3 | IFN-β | IFN-γ | IL-1β | IL-6 | IL-8 |
| 1 | mock | 15.285 | 0.669 | 6.761 | 22.01 | 5.93 | 0.97 | 6.47 | 16.61 | 4788.965 |
| 2 | mock | 29.836 | 1.182 | 11.246 | 48.61 | 19.131 | 1.7 | 14.5 | 29.59 | 8570.515 |
| 3 | mock | 41.735 | 1.31 | 13.626 | 52.74 | 18.155 | 2.1 | 15.45 | 36.05 | 11628.465 |
| 1 | LPS | 14.706 | 0.937 | 6.761 | 22.95 | 5.931 | 0.97 | 5.41 | 13.1 | 4280.483 |
| 2 | LPS | 20.446 | 0.862 | 8.663 | 26.73 | 6.212 | 0.97 | 10.08 | 19.67 | 6299.808 |
| 3 | LPS | 23.085 | 0.981 | 9.112 | 43.53 | 17.831 | 1.56 | 9.32 | 24.3 | 6747.016 |
| 1 | R848 | 18.828 | 0.894 | 9.553 | 35.17 | 10.894 | 1.32 | 13.06 | 27.03 | 6533.723 |
| 2 | R848 | 19.263 | 0.894 | 8.663 | 32.37 | 9.462 | 1.22 | 12.33 | 23.85 | 7074.38 |
| 3 | R848 | 18.62 | 0.729 | 8.205 | 22.01 | 7.629 | 0.97 | 9.07 | 21.63 | 5521.605 |
| 1 | CpG | 11.033 | 0.574 | 6.188 | 13.41 | 5.931 | 0.97 | 4.61 | 5.9 | 4872.379 |
| 2 | CpG | 13.423 | 0.81 | 8.205 | 23.9 | 5.931 | 0.97 | 8.17 | 9.58 | 6586.405 |
| 3 | CpG | 13.784 | 0.831 | 9.112 | 29.55 | 6.492 | 1.51 | 9.07 | 14.13 | 6489.428 |
| 1 | poly(I:C) | 8008.032 | 1.159 | 324.505 | 63.66 | 261.164 | 1.41 | 16.15 | 456.82 | 19457.155 |
| 2 | poly(I:C) | 9717.693 | 1.925 | 370.669 | 90.86 | 302.521 | 2.61 | 20.79 | 693.4 | 23680.885 |
| 3 | poly(I:C) | 9967.855 | 2.144 | 436.984 | 114.72 | 336.636 | 2.82 | 24.12 | 762.95 | 25097.665 |
| 1 | 3pRNA | 7364.434 | 1.047 | 140.25 | 61.85 | 241.609 | 1.41 | 18.45 | 48.65 | 9523.413 |
| 2 | 3pRNA | 7845.313 | 1.047 | 159.882 | 68.18 | 236.71 | 1.51 | 21.23 | 53.93 | 12032.822 |
| 3 | 3pRNA | 8424.087 | 1.345 | 167.729 | 77.14 | 264.936 | 1.51 | 19.8 | 61.12 | 11463.586 |
| 1 | c-GAMP | 133.378 | 1.024 | 16.947 | 44.92 | 28.268 | 1.36 | 10.96 | 64.01 | 5924.002 |
| 2 | c-GAMP | 102.066 | 0.64 | 10.832 | 22.95 | 5.931 | 0.97 | 8.29 | 54.51 | 5697.244 |
| 3 | c-GAMP | 115.437 | 1.069 | 12.456 | 36.1 | 16.54 | 1.41 | 13.3 | 70.44 | 6955.899 |

We next evaluated the response to viral PAMPs on phagocytes, which are overrepresented in the BALF. In particular, we activated bacterial or viral sensors in bulk peripheral blood mononuclear cells (PBMCs), or in monocytes or conventional dendritic cells (cDCs) isolated form PBMCs. Monocyte-derived DCs (moDCs) were used as a comparison. Regarding the production of IFNs, each cell population was found to respond differentially to PRR stimulation: while PBMCs were particularly able to produce IFN-II in response to viral and bacterial ligands, cDCs were uniquely capable of producing very high levels of IFN-λ2,3, and to a lesser extent of IFN-λ1, solely in response to TLR3 stimulation (**Figure 5C-F, Table 8**). This is in agreement with our previous data, which show that in the mouse system, cDCs are the major producers of IFN-III in response to activation of the TLR3/TRIF pathway (*23*). When these analyses were extended to NF-kB-dependent (IL-1β, IL-6, TNF-α, IL-8) as well as to type II cytokine (IL-10, GM-CSF), and chemokine (CXCL10) production, each cell type revealed a unique pattern of protein production (**Supplementary Figure 5D-G, Table 8**), underscoring the complexity and cell-specificity of the inflammatory response.

**TABLE 8.**
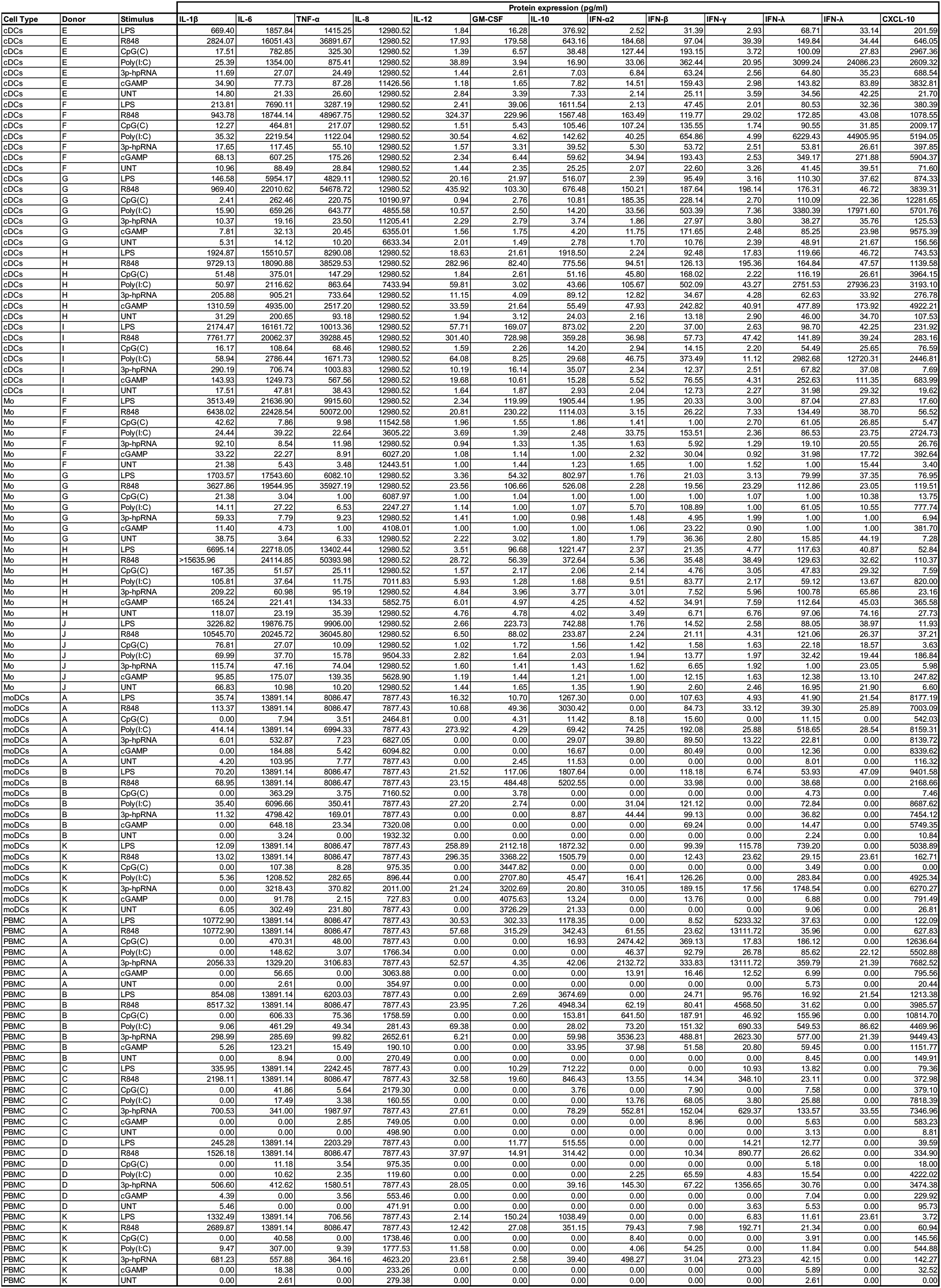
Cytokine protein production by blood-derived phagocytes stimulated with an array of PAMPs. Cytokine protein levels (pg/ml) produced by blood-derived human cDCs, monocytes, PBMCs, and moDCs stimulated with agonists of PRRs as described in **Figure 5**.

Collectively, these data demonstrate that epithelial cells preferentially produce IFN-λ1 upon RIG-I/MDA-5 or TLR3 stimulation, while TLR3 is the major driver of IFN-III production by cDCs. They also show that specific members of the IFN-I and IFN-III classes are differentially produced upon stimulation of PRRs on immune and epithelial cells.

## Discussion

COVID-19 has caused more than 2.5 million deaths globally, and has had devastating societal and economic effects. Notwithstanding the rising expectations linked to the rollout of the COVID-19 vaccines, a better understanding of the molecular underpinnings that drive the severe pathology in patients infected with the SARS-CoV-2 virus is imperative in order to implement effective additional prophylactic and/or therapeutic interventions. IFN-I and IFN-III are potent anti-viral cytokines and, among available therapeutic options, the potential of using clinical grade recombinant IFN-I or IFN-III as therapeutics has raised much hope and interest (*41*). However, a more complete understanding of how IFNs affect the development of COVID-19 is an essential first step for designing such IFN-related therapeutic interventions. To date, though, opposing evidence has complicated our view of the role played by members of the IFN-I and IFN-III families during SARS-CoV-2 infection. Here, we present analyses of respiratory samples of a cohort of over 200 SARS-CoV-2 patients with different clinical manifestations of COVID-19. Similar analyses were also performed on respiratory samples from patients with non-SARS-CoV-2 ARDS, as well as with other non-infectious clinical conditions. We scrutinized gene and protein expression along the respiratory tract in these individuals and found higher levels of IFNs in the lower airways of individuals with severe COVID-19 compared to other ARDS patients. In particular, IFN-III production was most exuberant in the severely ill patients. Of note, we found no correlation between cytokine levels in the lungs and in the plasma. In contrast to the lower airways, an efficient production of IFN-III, and to a lesser extent of IFN-I, was detected in the upper airways of patients with a high viral load but lower risk of developing severe COVID-19. In particular, individuals younger than 70 years of age, who are known to be less susceptible to severe COVID-19 and COVID-19-related death, showed a stronger correlation between IFN levels and viral load. In keeping with a protective effect of the IFNs in the upper airways, we found that patients with mild manifestations exhibited elevated levels of IFN-III at high viral loads, while hospitalized patients didn’t show this correlation. And finally, we observed that different cell types produce either IFN-I or IFN-III in response to specific microbial ligands. Together, these data suggest that the variability in levels of specific members of the IFN families along the respiratory tract, and also among different groups of patients, may be at least partially explained by the cell types that are enriched and/or activated under the various conditions. Our results establish that the differences in the production of inter and intra-family IFN members are caused by a dynamic process that varies based on the localization of the response, and suggest that the IFNs play opposing roles at distinct anatomical sites, either preventing or facilitating the severity of COVID-19. These findings will be fundamental for designing appropriate pharmacological interventions to prevent infection with SARS-CoV-2 or for dampening the severity of COVID-19.

We and others have recently demonstrated that IFN-IIIs, long believed to be initiators of anti-viral responses at mucosal surfaces that are less detrimental than those induced by type I IFNs (*14*), also profoundly alter lung barrier function during a viral infection (*23, 24*). Our recent report describes that SARS-CoV-2 induced production of IFNs in the lower airways of a small cohort of 10 patients with COVID-19 and 5 healthy controls. Here, we have extended this analysis, and partially revised our conclusions, by showing not only that severely affected COVID-19 patients are characterized by the highest levels of IFNs (at the mRNA as well as protein levels), but also that high levels of SARS-CoV-2 induce the efficient production of IFN-III in the upper airways. Moreover, IFN production is increased in the upper airways of younger patients, and is associated with milder disease manifestations. These data support the hypothesis that IFNs have opposing roles along the respiratory tract, and reconcile some of the seemingly contradictory findings on IFNs in COVID-19 patients. Efficient initiation of IFN production in the upper airways (in younger and mildly infected patients) can lead to a more rapid elimination of the virus and may limit viral spread to the lower airways. The report that 15% of severe COVID-19 patients are characterized by defects in IFN signaling strongly supports the hypothesis that the early production of IFNs in the upper airways protects against SARS-CoV-2, and that in the absence of such production, SARS-CoV-2 may spread to the lower respiratory tract and cause a more severe illness. On the other hand, when the virus escapes immune control in the upper airways, the IFN production that is potently boosted in the lungs likely contributes to the cytokine storm and associated tissue damage that is typical of severe COVID-19. We also found that IFN-γ, which reportedly aggravates the severity of COVID-19 independently of viral the load, is elevated in the BALF of severe patients. However, since none of the IFNs tested in this study showed a positive correlation between local production in lung cells and systemic levels in the plasma, inferring the inflammatory landscape in the lungs based on plasma measurements remains a challenge.

Another novel finding in the present study is that the type of IFN produced in response to different PRR pathways varies according to cell type. This may explain the varying pattern of the expression of IFNs along the respiratory tract. Epithelial cells of the upper and lower respiratory tracts express viral entry receptors, which facilitate the entrance and replication of SARS-CoV-2 in these cells (*32–35*). Although the capacity of these cells to recognize and respond to viral components is confounded by the presence of SARS-CoV-2 effector proteins (which block immune recognition and IFN production) (*42–45*), our finding that the high viral load in the upper airways induces a potent immune response is compatible with reports from other groups that show IFN induction in epithelial cells upon SARS-CoV-2 infection (*28*). Note that viral loads in our cohort were not correlated with disease severity, supporting the view that a high viral load in the upper airways favors the production of IFNs, which in turn serve a protective role. This is in keeping with the recent observation that epithelial cell-intrinsic IFN-dependent responses limit viral spread, and that SARS-CoV-2 induces a more severe pathology in the absence of such responses (*35*). Moreover, there is a growing consensus that a high viral load in the respiratory tract is not *per se* a prognostic marker of clinical severity (*46, 47*). High viral loads in the upper airways may therefore be associated to a protective immune response in young individuals, whilst eliciting a dysregulated inflammatory response in older patients, as observed in our study.

Our findings shed new light on the nature of the IFNs and on the molecular pathways that drive intrinsic immunity. In particular, we have shown that IFN-III are highly represented in the upper airways of younger and/or less severely infected patients, and that epithelial cells preferentially produce this type of IFN. Although additional studies are needed to directly link specific IFNs to particular cell types, our data support the possibility that efficient activation of epithelial cells in the upper airways, and induction of the potent IFN-III response, protect against severe COVID-19.

In agreement with our previous findings (*23*), we show that, cDCs are major producers of IFN-III, and especially of IFN-λ2,3. These IFNs are characteristically expressed along the lower airways of severe COVID-19 patients. Epithelial cells can be directly infected by SARS-CoV-2, but phagocytes are believed to host only abortive infections (*33, 35, 36*). Most immune cells respond to SARS-CoV-2 upon recognizing either the virus or host-derived molecules (*40*). Indeed, we found that IFN-III production by phagocytes is driven by the TLR3 pathway, in humans as well as in mice. TLR3 recognizes double-stranded intermediates of SARS-CoV-2 replication that may be released by bystander-infected cells (*48*). Thus, while phagocyte-derived IFN-λ2,3 may contribute to raise IFN-III levels produced by epithelial cells, it may also drive the detrimental effects of IFN-III in the lower airways. Indeed, the lower airways are where extensive cell death during severe COVID-19 can initiate a negative feed-back loop that favors tissue damage, along with the release of viral intermediates and host inflammatory ligands that further exacerbate IFN production.

Finally, our findings highlight the importance of the timing of production and/or administration of IFNs during COVID-19. It has been proposed (*49*) and experimentally proven (*45, 50, 51*) that early administration of IFNs may prevent infection and/or favor SARS-CoV-2 clearance, consistent with previous mouse studies on highly pathogenic human coronaviruses (*52*). A retrospective study on the use of recombinant (r)IFN-I corroborates these findings (*53*). Yet the results of clinical trials that employed clinical grade human rIFN-λ were inconclusive: early administration of IFN-III in SARS-CoV-2-positive patients, ranging between mild and asymptomatic manifestations, did not shorten the time of hospitalization (*54*). In contrast, another study documented that if IFN-IIIs are dispensed early after symptom onset or to asymptomatic patients, this induces a faster decline in viral load and faster remission in patients with an elevated viral load (*55*). Our data are in agreement with the idea that early administration (before infection or early after symptom onset) of exogenous IFN-IIIs may be an effective therapeutic intervention, and that targeting the upper airways, while avoiding systemic administration as previously proposed (*49*), represents the best way to exploit the anti-viral activities of IFNs.

In conclusion, our data define the anatomical map of inter and intra-family production of IFNs, and highlight how IFN production is linked to the different outcomes of COVID-19, based on the location of the IFN response. Our findings reconcile a large portion of the literature on IFNs, and further stress the key role played by IFN-III, compared to IFN-I, at mucosal surfaces during life-threatening viral infections.

## Acknowledgments

We thank the members of the Zanoni and Mancini laboratories for thoughtful discussion and comments on the project. IZ is supported by NIH grants 1R01AI121066, 1R01DK115217, NIAID-DAIT-NIHAI201700100 and holds an Investigators in the Pathogenesis of Infectious Disease Award from the Burroughs Wellcome Fund. Research done in the Wack lab was funded in whole, or in part, by the Wellcome Trust [FC001206]. For the purpose of Open Access, the author has applied a CC BY public copyright license to any Author Accepted Manuscript version arising from this submission. S.C. and A.W. were supported by the Francis Crick Institute which receives its core funding from Cancer Research UK (FC001206), the UK Medical Research Council (FC001206) and the Wellcome Trust (FC001206). We thank Renato Ostuni and the Ostuni laboratory at TIGET, Hospital San Raffaele, Italy, for granting access to the ViiA7 Real-Time PCR System. LP and FM thank Drs. Daniele Lilleri and Chiara Fornara, Laboratories of Genetics, Transplantology and Cardiovascular Diseases, and Biotechnology Laboratories, IRCCS Policlinico San Matteo Foundation, Pavia, Italy, for granting the use of the cytofluorimeter.

## Author Contributions

BS and AB (Achille Broggi) designed, performed, analyzed the experiments and edited the text; LP, VF, LS, AB (Andrea Bottazzi), TF, CR, FM performed the protein analyses on the BALF fluids; SC and AW designed and performed the analyses on human lung epithelial cells; AA, EL, ET and AEP helped with statistical analyses; RF, SS, NC, MC, LM, FAF helped in the analysis of the gene quantification from the swabs; NM designed the experiments, supervised the project and edited the text; IZ conceived and supervised the project, designed the experiments and wrote the paper.

## Materials and Methods

### Reagents and antibodies

For in vitro studies, we used LPS (ALX-581-013-L002) from ENZO, Poly (I:C) HMW (tlr-pic), R848 (tlr-r848), CpG(C) (tlrl-2395), 2’3’cGAMP (tlrl-nacga23-02) and 3p-hpRNA/LyoVec (tlrl-hprnalv) purchased from Invivogen. Lipofectamine 3000 Transfection Reagent (L3000-008) was purchased from Invitrogen. Phagocytes were cultured in RPMI 1640 medium + GlutaMAX (72400-047) with 1% Penicillin-Streptomycin (15140122) purchased from Thermo Fisher, and 10% fetal bovine serum (FBS). HBECs were cultured as previously described (Major et al., 2020). For flow cytometry we used PerCP/Cy5.5 CD14 (clone HCD14), APC/Cyanine7 HLA-DR (clone L243), PE/Cy7 CD11c (clone 3.9) and PE CD141 (clone M80) antibodies purchased from Biolegend.

### Clinical samples for gene expression analysis

Nasopharyngeal swabs were collected using FLOQSwabs® (COPAN) in UTM® Universal Transport Medium (COPAN) from 152 SARS-CoV-2 positive patients and from 20 negative subjects undergoing screening for suspected social contacts with SARS-CoV-2 positive subjects. Nasopharyngeal swabs were collected at San Raffaele Hospital (Milan, Italy) from April to December 2020. BALF was obtained from severe SARS-CoV-2-positive patients hospitalized at San Raffaele Hospital (Milan, Italy) from March to May 2020. See **Table 1**, **Table 4**, **Table 5** for patient information. All samples were stored at −80°C until processing. 500 μl of each BALF and swab sample were lysed and used for RNA extraction (see *RNA extraction protocol and Real-Time PCR for clinical samples and HBECs*).

Data were obtained from the COVID-BioB clinical database of the IRCCS San Raffaele Hospital. The study was approved by the Ethics Committee of San Raffaele Hospital (protocol No. 34/int/2020) and was registered on ClinicalTrials.gov (NCT04318366). All patients signed an informed consent form. Our research was in compliance to the Declaration of Helsinki.

### Evaluation of SARS-CoV-2 RNA amount in clinical samples

The viral load was inferred on nasopharyngeal swabs through cycle threshold (Ct) determination with Cobas® SARS-CoV-2 Test (Roche), a real-time PCR dual assay targeting ORF-1a/b and E-gene regions on SARS-CoV- 2 genome. The mean between ORF-1a/b and E Ct was used as an indirect measure of the viral load. Non-infectious plasmid DNA containing a specific SARS-CoV-2 sequence and a pan-Sarbecovirus sequence are used in the test as positive control. A non-Sarbecovirus related RNA construct is used as internal control. The test is designed to be performed on the automated Cobas® 6800 Systems under Emergency Use Authorization (EUA). The test is available as a CE-IVD test for countries accepting the CE-mark.

### Clinical samples for cytokine quantification in BALF and plasma

BALF from 30 SARS-CoV-2 positive patients hospitalized in the Intensive Care Unit (ICU) at Luigi Sacco Hospital (Milan, Italy) were collected from September to November 2020. Blood from 17 of these patients was also collected on the same day. BALF from patients affected by ARDS (9 in total, 5 of which were diagnosed H1N1 influenza A virus) were collected from February 2014 to March 2018. Samples from: lung fibrosis patients (10) were collected from May 2018 to September 2020; sarcoidosis patients (10) were collected from August to July 2020; lung transplant patients (10) were collected from January 2018 to September 2020 by IRCCS Policlinico San Matteo Foundation (Pavia, Italy). None of the patients affected by lung fibrosis, sarcoidosis or that received lung transplant was diagnosed a respiratory viral or bacterial infection. See **Table 2** for patient information. BALF specimens from COVID-19 patients were managed in a biosafety level 3 laboratory until viral inactivation with a 0.2% SDS and 0.1% Tween-20 solution and heating at 65 °C for 15 min. Cell-free BALF supernatants were stored at − 20 °C until analysis. Blood was centrifuged at 400g for 10 minutes without brake and plasma was stored at − 20 °C until analysis. Samples were processed with LEGENDplex^TM^ (740390, BioLegend) according to manufacturer’s instructions and read by flow cytometry.

Research and data collection protocols were approved by the Institutional Review Boards (Comitato Etico di Area 1) (prot. 20100005334) and by IRCCS Policlinico San Matteo Foundation Hospital (prot. 20200046007). All patients signed an informed consent form. Our research was in compliance to the Declaration of Helsinki.

### Culture of primary HBECs and in vitro stimulation

Primary human bronchial epithelial cells were purchased from Lonza and cultured as per manufacturer’s instructions. In brief, cells were expanded in a T-75 flask to 60% confluence and then trypsinized and seeded (3×10^4^ cells/transwell) onto 0.4 μm pore size clear polyester membranes (Greiner) coated with a collagen solution. Cells were grown in submersion until confluent, and then exposed to air to establish an air-liquid interface (ALI). At ALI day 15, cells were stimulated with LPS (100 ng/ml), R848 (10 μg/ml), CpG(C) (1 μM), Poly (I:C) (50 μg/ml), Poly (I:C) (1 μg/10^6^ cells) + Lipofectamine, 3p-hpRNA/LyoVec (100 ng/ml), and cGAMP (10 μg/ml). Supernatants and cell lysates were collected 24 hours post treatment. Supernatants were processed with LEGENDplex^TM^ (740390, BioLegend) according to manufacturer’s instructions and read by flow cytometry. Lysates were processed for RNA extraction as described below.

### Isolation of human phagocytes and in vitro stimulation

Human phagocytes were isolated from collars of blood received from Boston Children’s Hospital blood donor center. Briefly, blood was diluted 1:2 in PBS and PBMCs were isolated using a Histopaque (1077-1, Sigma) gradient. Monocytes were positively selected from PBMCs with CD14 MicroBeads (130-050-201, Miltenyi Biotec) by MACS technology. MoDCs were differentiated from monocytes in the presence of GM-CSF 20ng/ml (PeproTech, 300-03) and IL-4 20ng/ml (PeproTech 200-04) for 7 days. MoDCs differentiation was tested for CD14 downregulation and HLA-DR expression (**Figure Supplementary 6A, B**). cDCs were positively selected from PBMCs with CD141 (BDCA-3) MicroBead Kit (130-090-512, Miltenyi Biotec) by MACS technology (**Figure Supplementary 6C**). Cells were stimulated with LPS (100 ng/ml), R848 (10 μg/ml), CpG(C) (1 μM), Poly (I:C) (50 μg/ml), 3p-hpRNA/LyoVec (2.5 μg/ml), and cGAMP (10 μg/ml). Supernatants were collected 24 hours post treatment and stored at − 20 °C until analysis. Supernatants were processed with LEGENDplex^TM^ (740390, BioLegend) according to manufacturer’s instructions and read by flow cytometry.

### RNA extraction protocol and Real-Time PCR from clinical samples and HBECs

RNA was extracted from nasopharyngeal swabs, BALFs and HBECs culture lysates using Pure Link RNA Micro Scale kit (12183016, Thermo Fisher) according to manufacturer’s instruction, including in-column DNase treatment. Reverse transcription was performed using SuperScript^TM^ III First-Strand Synthesis System (18080051, Invitrogen) according to manufacturer’s instruction. qRT-PCR analysis was then carried out with Taqman^TM^ Fast Advanced Master Mix (4444963) by using specific Taqman^TM^ Gene Expression Assays from Thermo Fisher. *IFNL1 (*Hs01050642_gH)*, IFNL2,3 (*Hs04193047_gH)*, IFNL4* (Hs04400217_g1), *IFNB1* (Hs01077958_s1), *IFNA2* (Hs00265051_s1)*, IFNA4* (Hs01681284_sH)*, IL1B* (Hs01555410_m1) and *IL6* (Hs00174131_m1) expression was assessed with respect to the housekeeping gene *GAPDH* (Hs99999905_m1). All transcripts were tested in triplicate for each sample on ViiA7 Real-Time PCR System (Thermo Fisher) for clinical samples and on Quantastudio 3 Real-Time PCR System (Thermo Fisher) for HBECs.

### Statistical Analyses

One-way ANOVA with Turkey’s post-hoc test was used to compare continuous variables among multiple groups. Kruskal-Wallis test with Dunn’s post-hoc test or Multiple Mann-Whitney tests with Holm-Šídák method were used instead when data did not meet the normality assumption. Fisher’s exact test was used to compare categorical variables. Spearman correlation analysis was used to examine the degree of association between two continuous variables. To establish the appropriate test, normal distribution and variance similarity were assessed with the D’Agostino-Pearson omnibus normality test.

Cluster analysis with unbiased K-mean methods based on the expression of IFN-I, IFN-III and the proinflammatory cytokine IL-1β were used to classify a subset of COVID-19 patients into 3 exclusive clusters. Heatmaps and K-mean clustering were generated in R and visualized with the ComplexHeatmap package. Clustering analysis was performed using Euclidean distances. Estimated (K) value was selected based on the elbow point cluster number. Logistic regression models were performed to estimate the association of gene expression as binary outcome within viral load terciles (defined by mean viral RNA CT <20, >20 and <30, > 30), and clusters (cluster 1, cluster 2 and cluster 3). Interaction between viral load terciles and age groups (≥70 years vs <70 years) were tested to detect significant difference between elder patients and young patients in their gene expression response to different levels of viral load. All statistical analyses were two-sided and performed using Prism9 (Graphpad) software or SAS version 9.4 (SAS Institute).

## SUPPLEMENTARY INFORMATION

### SUPPLEMENTARY FIGURES AND FIGURE LEGENDS

**SUPPLEMENTARY FIGURE 1.**
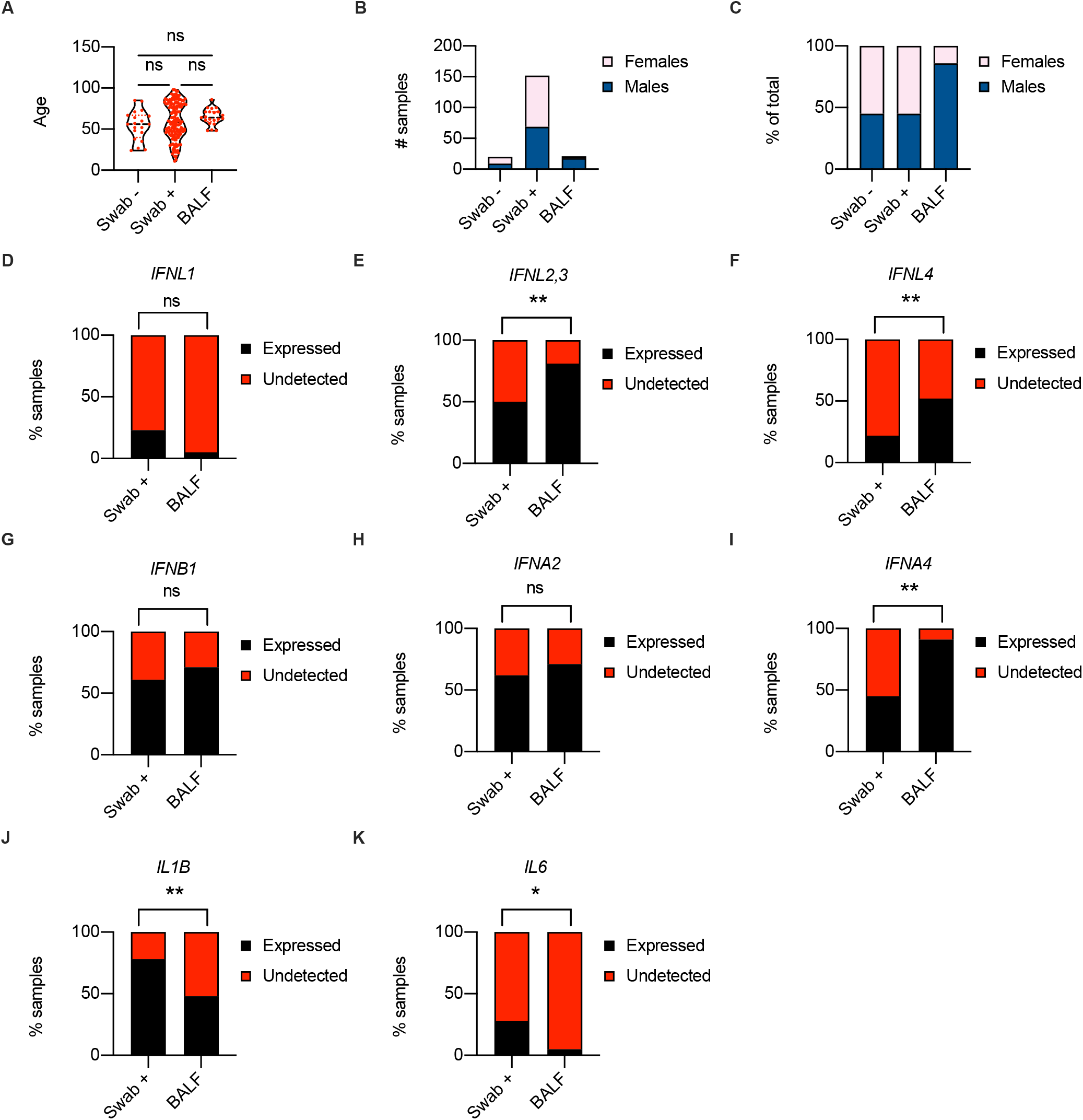
(**A-C**) Age distribution (**A**), number (**B**) and percentage (**C**) of females and males in cohorts of patients (Swab −, Swab + and BALF) analyzed in **Figure 1A-H**. (**D-K**) Percentage of patients that express (Expressed, black bars) or not (Undetected, red bars) *IFNL1* (**D***), IFNL2,3* (**E***), IFNL4* (**F**), *IFNB1* (**G**), *IFNA2* (**H**), *IFNA4* (**I**), *IL1B* (**J**), and *IL6* (**K**) in SARS-CoV-2^+^ swabs (Swab +) and in the BALF from SARS-CoV-2^+^ patients (BALF). Statistics: (**A**) One-way ANOVA test with Turkey’s post-hoc test: ns, not significant (*P*>0.05); \**P*<0.05, \*\**P*<0.01, \*\*\**P*<0.001, and \*\*\*\**P*<0.0001. (**D-K**) Fisher’s exact test; ns, not significant (*P*>0.05); \**P*<0.05, \*\**P*<0.01, \*\*\**P*<0.001, and \*\*\*\**P*<0.0001.

**SUPPLEMENTARY FIGURE 2.**
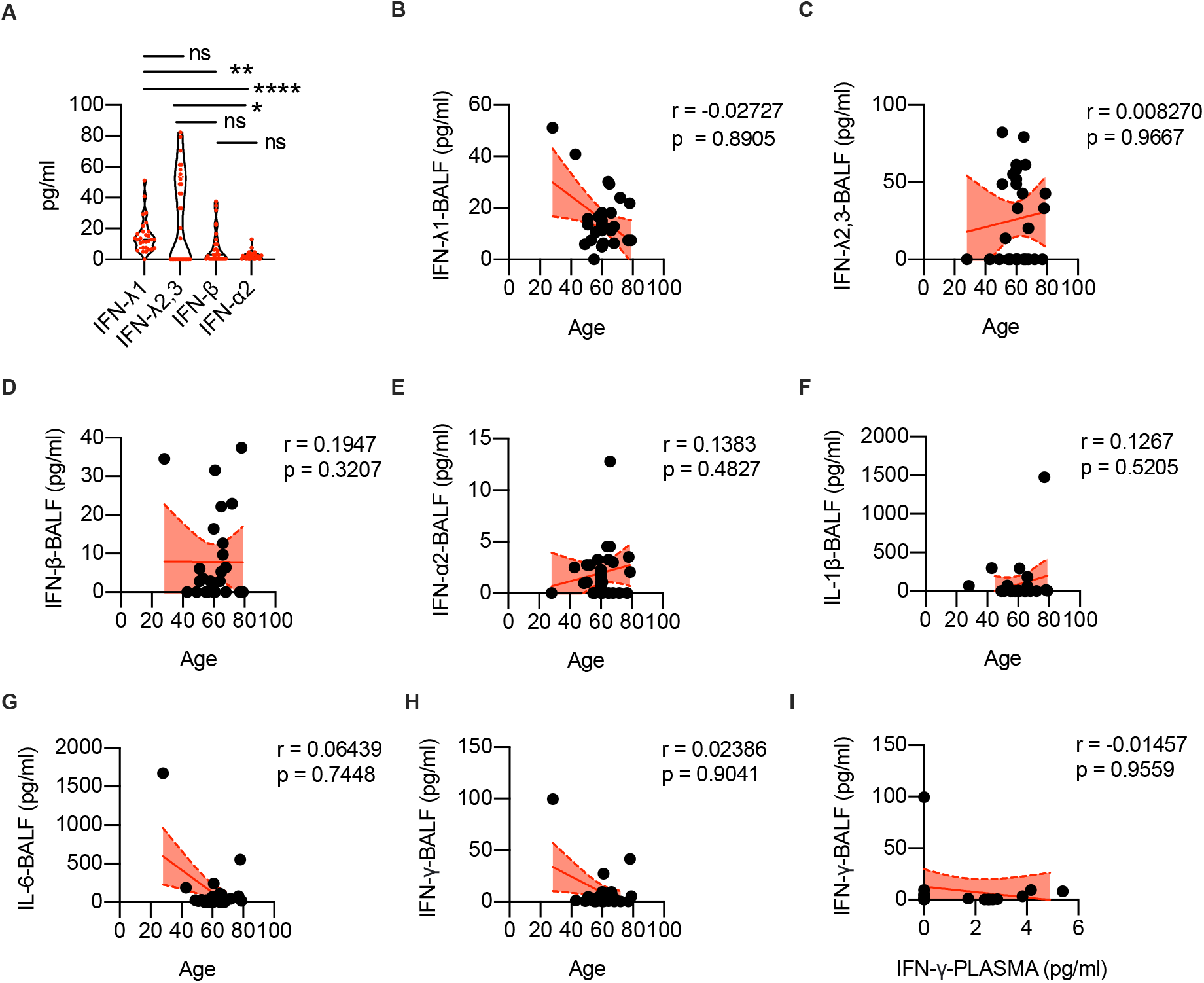
(**A**) Protein levels of IFN-λ1, IFN-λ2,3, IFN-β, IFN-α2 measured in the BALF of COVID-19 patients. Each dot represents a patient. Violin plots are depicted. (**B-H**) IFN-λ1 (**B**), IFN-λ2,3 (**C**), IFN-β (**D**), IFN-α2 (**E**), IL-1β (**F**), IL-6 (**G**) and IFN-γ (**H**) protein levels in the BALF of COVID-19 patients are plotted over age. (**I**) IFN-γ protein levels in the BALF are plotted against protein levels in the plasma of each COVID-19 patient. (**B-I**) Each dot represents a patient. Linear regression lines (continuous line) and 95% confidence interval (dashed line and shaded area) are depicted in red. Spearman correlation coefficients (r) and p-value (p) are indicated. Statistics: (**A**) Kruskal-Wallis test with Dunn’s post-hoc test: ns, not significant (*P*>0.05); \**P*<0.05, \*\**P*<0.01, \*\*\**P*<0.001, and \*\*\*\**P*<0.0001.

**SUPPLEMENTARY FIGURE 3.**
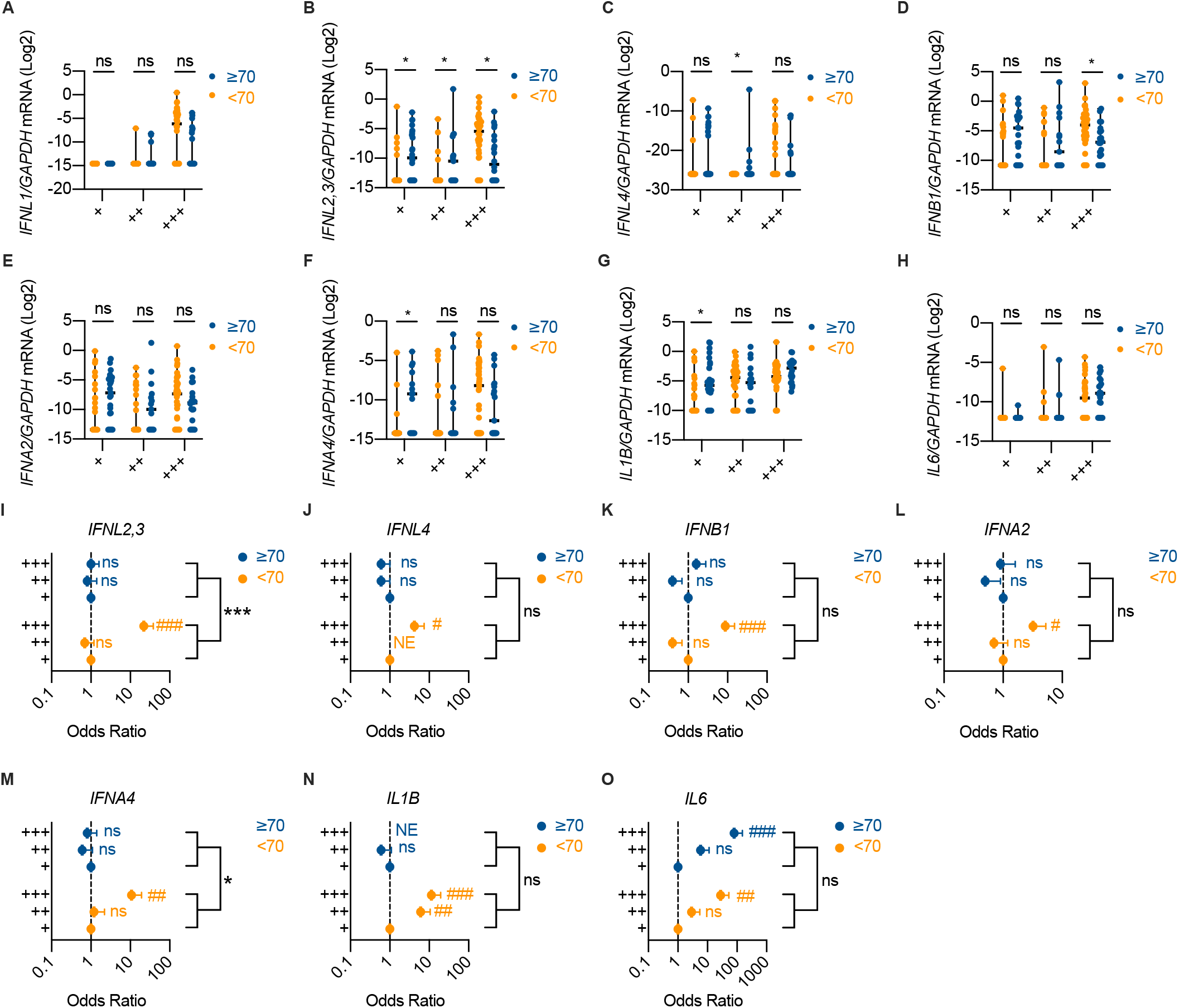
(**A-H**) *IFNL1* (**A***), IFNL2,3* (**B***), IFNL4* (**C**), *IFNB1* (**D**), *IFNA2* (**E**), *IFNA4* (**F**), *IL1B* (**G**), and *IL6* (**H**) mRNA expression in SARS-CoV-2^+^ swabs divided in viral load terciles (“+++”, “++”, “+”) defined by mean viral RNA CT (<20, >20 and <30, > 30) in patients over 70 years old (≥70, blue dots and lines) and below 70 years old (< 70, orange dots and lines). Expression is plotted as log2 (*gene*/*GAPDH* mRNA *+* 0.5 x gene-specific minimum). Each dot represents a patient. Median with range is depicted. (**I-O**) Odds ratio of expressing *IFNL2,3* (**I***), IFNL4* (**J**), *IFNB1* (**K**), *IFNA2* (**L**), *IFNA4* (**M**), *IL1B* (**N**), and *IL6* (**O**) mRNA in “+++” and “++” with respect to “+” SARS-CoV-2^+^ swabs in ≥70 (blue dots and lines) and < 70 (orange dots and lines) patients. Symbols represent the odds ratio. Error bars represent the 95% confidence interval associated to the odds ratio. NE: not estimable, J) no patient in group expresses *IFNL4*, N) all patients in group express *IL1B*. Statistics: (**A-H**) Multiple Mann-Whitney tests with Holm-Šídák method: ns, not significant (*P*>0.05); \**P*<0.05, \*\**P*<0.01, \*\*\**P*<0.001, and \*\*\*\**P*<0.0001. (**I-O**) Odds ratio: ns, not significant (*P*>0.05); #*P*<0.05, ##*P*<0.01, ###*P*<0.001. Interaction analysis: ns, not significant (*P*>0.05); \**P*<0.05, \*\**P*<0.01, \*\*\**P*<0.001.

**SUPPLEMENTARY FIGURE 4.**
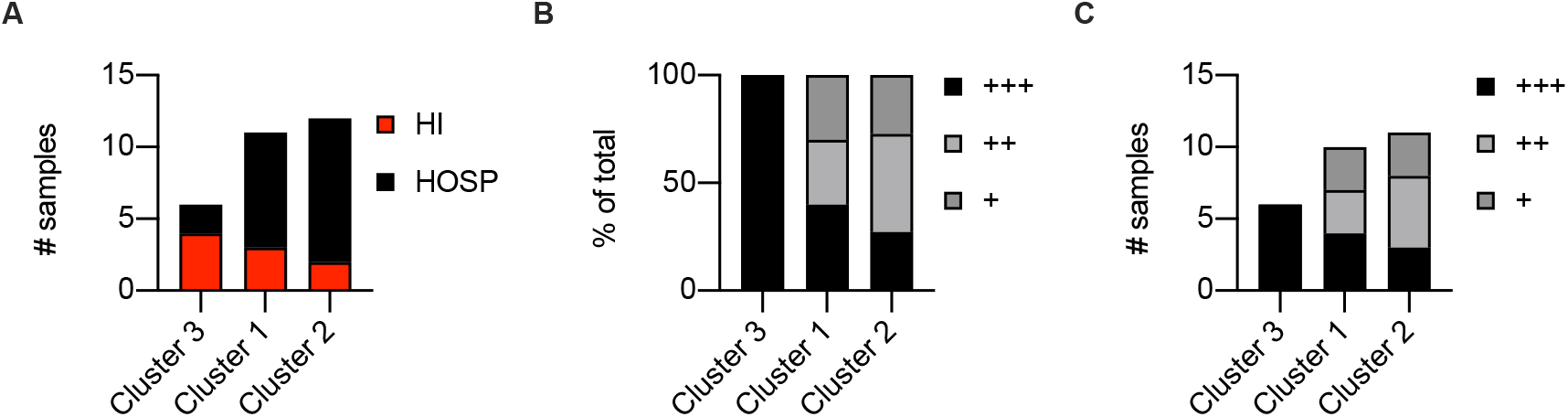
(**A**) Number of samples from each disease severity group (HOSP=hospitalized and HI=home-isolated) within each cluster identified in Figure 4I. (**B-C**) Percentage (**B**) and number (**C**) of samples from each viral load tercile (“+++”, “++”, “+”) within each cluster identified in Figure 4I. Viral load terciles (“+++”, “++”, “+”) are defined by mean viral RNA CT (<20, >20 and <30, > 30).

**SUPPLEMENTARY FIGURE 5.**
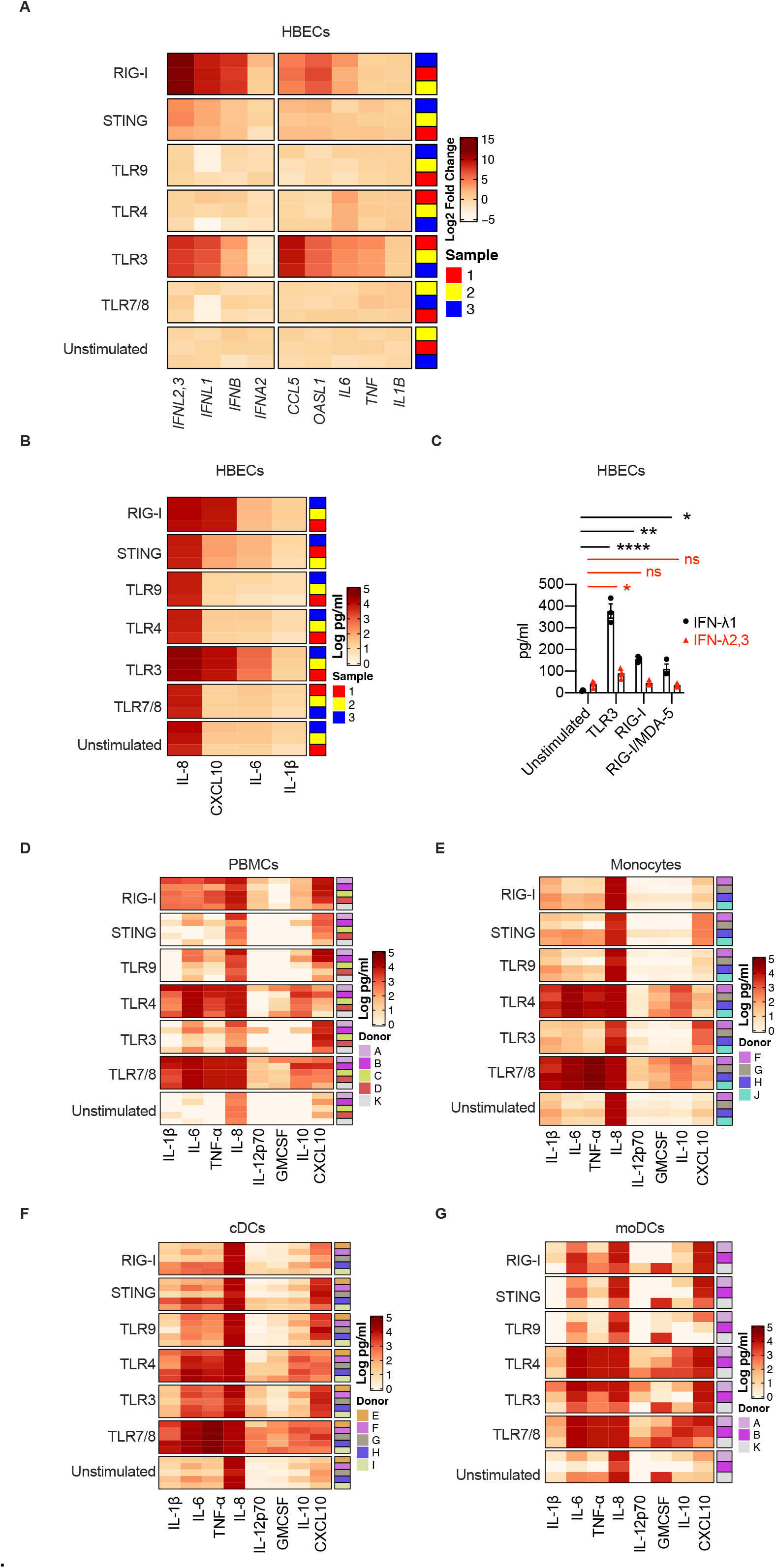
(**A-B**) HBECs were treated with 3p-hpRNA/LyoVec, cGAMP, CpG(C), LPS, Poly (I:C) and R848 for stimulation of RIG-I, STING, TLR9, TLR4, TLR3 and TLR7/8 respectively. **(A)** Heatmap representation of *IFNL2,3*, *IFNL1, IFNB1, IFNA2, CCL5, OASL1, IL6, TNF* and *IL1B* mRNA expression 24 hours after treatment. The color is proportional to Log2 (Fold Change) of each gene. Rows in each group represent biological replicates distributed as indicated in the legend. (**B**) Heatmap representation of IL-8, CXCL10, IL-6 and IL-1β production 24 hours after stimulation. The color is proportional to the Log10 transformed concentration (pg/ml) of each cytokine. Rows in each group represent a biological replicate. (**C**) IFN-λ1 and IFN-λ2,3 production by HBECs treated for 24h with PRR ligands. Poly (I:C) (TLR3), 3p-hpRNA/LyoVec (RIG-I) and transfected Poly (I:C) (RIG-I/MDA5) were used. Mean with SEM is depicted. Each dot represents a biological replicate. (**D-G**) Heatmap representation of IL-1β, IL-6, TNF-α, IL-8, IL-12p70, GMCSF, IL-10 and CXCL10 production by PMBCs (**D**), Monocytes (**E**), cDCs (**F**), moDCs (**G**) treated for 24h with PRR ligands. 3p-hpRNA/LyoVec (RIG-I), cGAMP (STING), CpG(C) (TLR9), LPS (TLR4), Poly (I:C) (TLR3), R848 (TLR7/8) were used. (**D-G**) The color is proportional to the Log10 transformed concentration (pg/ml) of each cytokine. Rows in each group represent different donors as depicted in the annotation on the right. Statistics: (**C**) One-Way ANOVA with Dunnett’s post-hoc test: ns, not significant (*P*>0.05); \**P*<0.05, \*\**P*<0.01, \*\*\**P*<0.001, and \*\*\*\**P*<0.0001.

**SUPPLEMENTARY FIGURE 6.**
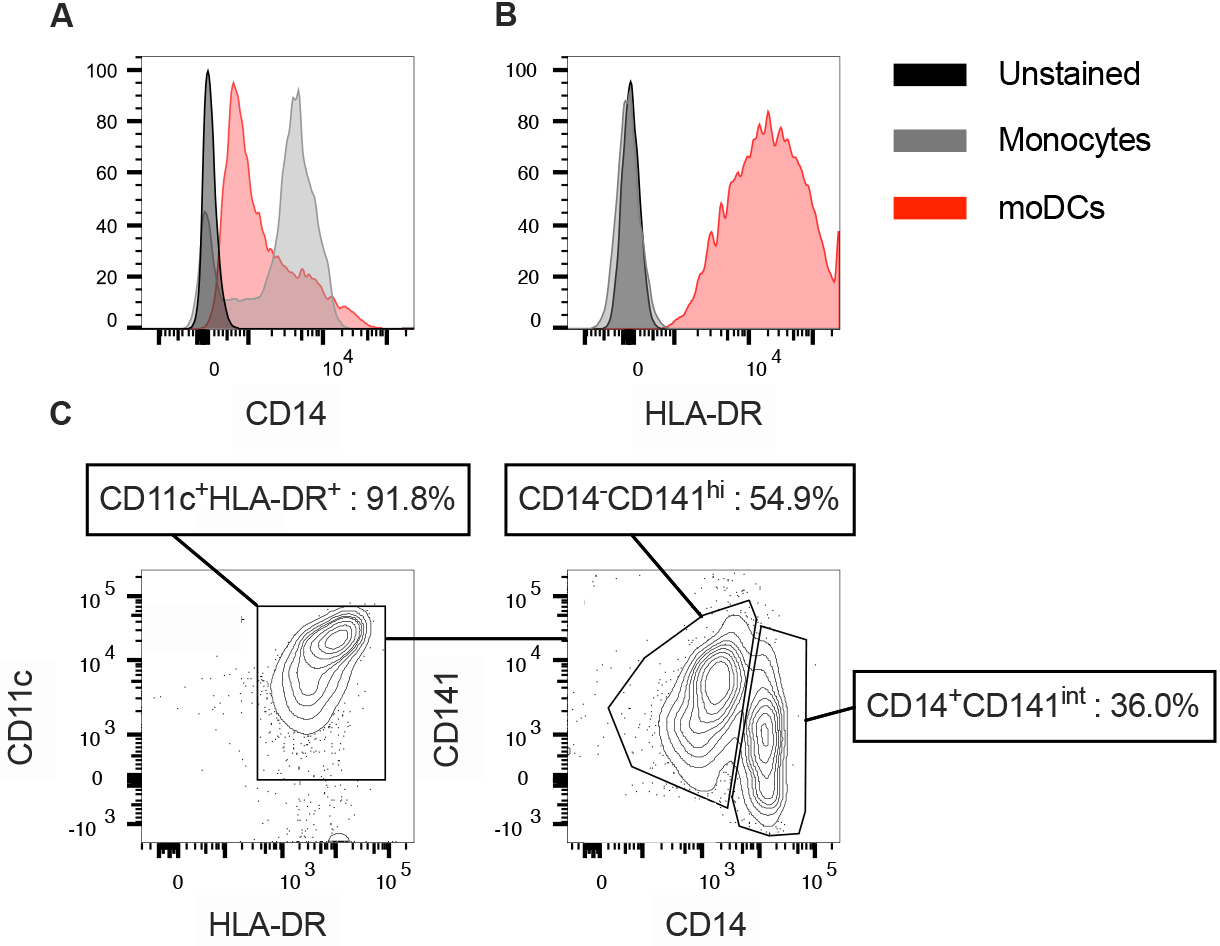
(**A-B**) CD14 (**A**) and HLA-DR (**B**) expression in blood monocytes and moDCs derived from blood monocytes was assessed by flow cytometry. **(C)** Representative flow cytometry plot of blood-derived cDCs. Percentage of total cDCs (CD11c^+^ HLA-DR^+^, left panel) or of CD141^hi^CD14^−^ and CD141^int^CD14^+^ DCs (right panel) are indicated.

